# Machine learning on magnetoencephalography data yields generalizable low-dimensional neural fingerprints that distinguish individuals across task conditions

**DOI:** 10.64898/2026.06.26.734424

**Authors:** Joonas Karhula, Anttoni Ojanperä, Ersin Yılmaz, Susanne Merz, Samuel Kaski, Riitta Salmelin

## Abstract

Individual brains are unique in structure and function. Functional differences are captured by neural fingerprints, which reflect individual differences in behavior and cognition as well as group-level changes related to neurodegenerative diseases. Most research efforts so far have focused on fingerprints comprising full functional connectomes. However, the high dimensionality of the connectomes can increase computational load and impede performance of machine learning methods in potential applications. A low-dimensional alternative that retains individual features of the full connectomes would thus be beneficial. The present study employed latent-noise Bayesian Reduced Rank Regression (lnBRRR) to learn low-dimensional latent spaces that capture individual features in functional connectivity and power spectral density data derived from MEG recordings. LnBRRR performance was assessed with low training set sizes (N=20-44), and against principal component analysis and linear discriminant analysis. Model performance was also assessed with task data, and the solutions were compared across task conditions with cosine similarity to establish whether individual features are altered by different cognitive processes. LnBRRR captured generalizable individual patterns already at N=20 but N=30-35 was needed to reach optimal test accuracies and to prevent potential overfitting. The model also achieved comparable performance to the alternative models. Latent fingerprints derived from task data attained comparable performance to resting-state latent fingerprints, and lnBRRR solutions were shown to generalize across conditions. Additionally, the model solutions for power spectral density data were discovered to be notably similar, yet differently rotated, over task conditions, suggesting that similar patterns of individual features were captured by the model regardless of the task condition. Altogether, the present results highlight lnBRRR as a potential tool for neuroimaging data analysis and demonstrate that individual differences in power spectral density are largely intrinsic and unaffected by varying cognitive processes.

## 1 INTRODUCTION

All human brains are individual with variable structures and functional connections. Recent interest in inter-individual differences has led to inquiry into neural fingerprints, which comprise inter-areal functional connections, and whether these fingerprints also explain behavioral differences. Previous work has shown that neural fingerprints derived from fMRI, EEG, and MEG can differentiate individuals and families as well as explain behavioral variables (Rocca et al., 2014; Finn et al., 2015; Leppäaho et al., 2019; Demuru and Fraschini, 2020; Da Silva Castanheira et al., 2021; Sareen et al., 2021; Haakana et al., 2024). In addition, distinct neural fingerprints have been observed for healthy and patient groups of various neurodegenerative diseases (Vecchio et al., 2018; Pusil et al., 2019; Stampacchia et al., 2024; Da Silva Castanheira et al., 2024), highlighting the potential of neural fingerprints for improvement of individualized diagnostics.

The majority of previous electrophysiological studies on neural fingerprints have utilized either full functional connectomes or power spectral features as fingerprints and identified participants based on inter- and intraparticipant Pearson’s correlations. In this approach, correlations are computed between two sets of neural fingerprints extracted from the same individuals, and for each participant the identification is considered successful if the intraparticipant correlation exceeds the interparticipant correlations. Despite the successful identification results (Rocca et al., 2014; Demuru and Fraschini, 2020; Da Silva Castanheira et al., 2021; Sareen et al., 2021), the high dimensionality of the connectomes and power spectral features could complicate their use in further applications as high-dimensional data can hinder machine learning performance and requires substantial computational resources to be processed on a large scale.

The problems related to fingerprint dimensionality have been tackled with several approaches in the literature. Random projections have been successfully applied with fMRI connectomes to alleviate the dimensionality issue (Duong-Tran et al., 2024). Although this approach demonstrated that relatively few randomly chosen connections suffice for distinguishing individuals successfully, neurobiological interpretation of the resulting low-dimensional fingerprint is complicated as the features are chosen randomly. In contrast to traditional machine learning models that suffer from high dimensionality, complex models, such as convolutional neural networks, can effectively utilize neural fingerprints without dimension reduction (Wang et al., 2019). These models, however, are also burdened by the interpretability problem. It is noteworthy that interpretability is not necessarily important if the aim is solely to identify individuals or predict other phenomena based on the neural fingerprints. However, if neurobiological interpretations based on neural fingerprints are desired, models that provide a transparent mapping between the fingerprints and the model output should be preferred for dimension reduction and predictive modeling.

Latent-noise Bayesian Reduced Rank Regression (lnBRRR) provides a transparent alternative for the fingerprint dimensionality problem. LnBRRR learns a low-dimensional latent space based on the observed data and user-chosen independent variables (Gillberg et al., 2016). Thus, when individual identifiers are used as independent variables, lnBRRR finds latent components comprised of functional connectivity (or power spectral density) patterns that distinguish individuals. Furthermore, lnBRRR enables simultaneous use of multiple independent variables (Gillberg et al., 2016), which grants it more flexibility compared to simpler models such as linear discriminant analysis (LDA). The model solution can be interpreted relatively easily because the mapping between the observed data and the latent space is linear. LnBRRR also acknowledges presence of structured noise in the observed data and assumes that the latent space conveys effects of both the independent variables and structured noise, which enables modeling of weak effects (Gillberg et al., 2016). The capability to find meaningful effects among noise is a notable benefit while working with M/EEG signals that can have low signal-to-noise ratio. Previous studies have shown that lnBRRR can differentiate both individuals and siblings to a high degree with M/EEG functional connectivity and spectral power fingerprints (Leppäaho et al., 2019; Haakana et al., 2024; Heikkinen et al., 2026).

Although lnBRRR has been proven suitable for learning neural fingerprints, various questions regarding the model – and, more generally, of the nature of the neural fingerprints – remain unanswered. As lnBRRR is intended for use in neuroimaging research, model performance and generalizability to unseen data must be examined with sample sizes common in neuroimaging literature (N=20-40). If such sample sizes were sufficient for the model, it would provide a useful tool in various neuroimaging research frames due to its aforementioned flexibility. LnBRRR has also not been put to test against simpler dimension reduction methods. Previous studies have shown that neural fingerprints can identify individuals across task conditions (e.g., by using one resting-state and one working-memory fingerprint) which suggests that some individual features remain unchanged by varying cognitive processes (Finn et al., 2015; Gratton et al., 2018; Colenbier et al., 2023). Whether task data can be utilized for lnBRRR fingerprinting, and whether the obtained fingerprints generalize or share similar components across task conditions remains uncharted.

The present study aimed to systematically cover these open questions. Human Connectome Project (HCP) MEG data (Larson-Prior et al., 2013; Van Essen et al., 2013) was utilized as it comprises various task conditions and has already been successfully used in previous fingerprinting studies. First, technical questions regarding lnBRRR were addressed. Model generalizability and reproducibility with low sample sizes were examined with N=20-44. Additionally, lnBRRR performance was benchmarked against principal component analysis (PCA) and LDA. Subsequently, lnBRRR was used to investigate the individual features in task data. Data from working-memory and story-math tasks were used to extract task-based individual fingerprints and their identification performance was compared with resting-state fingerprints. To establish whether resting-state and task fingerprints share similar individual patterns, cross-conditional identification was performed and the similarity of model solutions across the task conditions was assessed.

## 2 MATERIALS AND METHODS

### 2.1 Dataset and data splits

The Human Connectome Project (HCP) preprocessed MEG dataset was used for the present study (Larson-Prior et al., 2013; Van Essen et al., 2013). Resting-state (N=87, duration=6 min), working memory task (N=74, duration=10 min), and story-math task (N=75, duration=7 min) recordings were used. The resting-state data was recorded in supine position with eyes open. The working memory task was an N-back task, and the story-math task comprised trials during which the subject first listened to either a short story or a mathematical problem and was then required to answer a question related to the heard segment. Three resting-state recordings and two recordings of each task were available. Two recordings from each condition were selected for the present study to enable performance comparison between conditions.

The data was split in two ways for the experiments. Data split 1: The resting-state dataset comprising 31 monozygotic and heterozygotic sibling pairs and 25 subjects without siblings (total N=87) was used to estimate model performance with N=20-44 and to assess the similarity of model solutions between training sets. The participants were divided into two subsets with 43 and 44 participants. From these two subsets, training sets of 20, 25, 30, and 35 participants were subsequently selected. Therefore, lnBRRR was trained on two separate training sets with each N and tested with both the training set and the test set (the subset of N=43/44 not used in training). This enabled the test set size to remain unchanged while the training set size was reduced. Siblings were always included in the same subset. Data split 2: The 68 subjects with MEG recordings from all task conditions were divided into five cross-validation (CV) folds sized either 14 (three folds) or 13 (two folds) to estimate identification accuracy on unseen data. Siblings were always included in the same fold. The data split 2 was also used for benchmarking lnBRRR against PCA and LDA as well as to test neural fingerprints extracted from task data. Data from all 68 subjects were used for training lnBRRR for model similarity assessment across task conditions.

### 2.2 Source localization

FieldTrip toolbox was utilized for MEG data source analysis (Oostenveld et al., 2011). Prior to source analysis, the data from all conditions was cropped to 195 s length. Source estimation was performed with LCMV beamformer (Van Veen et al., 1997), subsequently to which the source estimated signals were clustered with PCA to 116 regions of interest defined by AAL atlas (Tzourio-Mazoyer et al., 2002). HCP data contains standardized head and source models for all participants, which were used for the beamformer. Story-math task data was split into story and math epochs, which were further processed independently, before source localization.

### 2.3 Functional connectivity

Based on results by Haakana et al. (2024), phase-locking value (PLV) (Lachaux et al., 1999), imaginary phase-locking value (iPLV) (Bruña et al., 2018), phase linearity measurement (PLM) (Baselice et al., 2019), and amplitude envelope correlation (AEC) (O’Neill et al., 2015) were selected for the present study. For functional connectivity calculations, the ROI signals were segmented to 45 non-overlapping four-second epochs (total length=180 s) with two seconds of real data padding on both sides. Following the segmentation the epochs were filtered to one of the frequency bands (Delta: 2-4 Hz, Theta: 4-8 Hz, Alpha: 8-13 Hz, Low beta: 13-20 Hz, High beta: 20-30 Hz, Gamma: 30-45 Hz) and Hilbert filtered, after which the padding was removed and functional connectivity calculated. Functional connectivity values were averaged over epochs to grant one value for each ROI pair (6670 pairs in total). Connectivity values for ROIs located in deep subcortical areas, cerebellum, and near the eyes were excluded to enhance reliability of the acquired metrics (Fig. 1). After exclusion, 1891 FC features remained.

**Figure 1.**
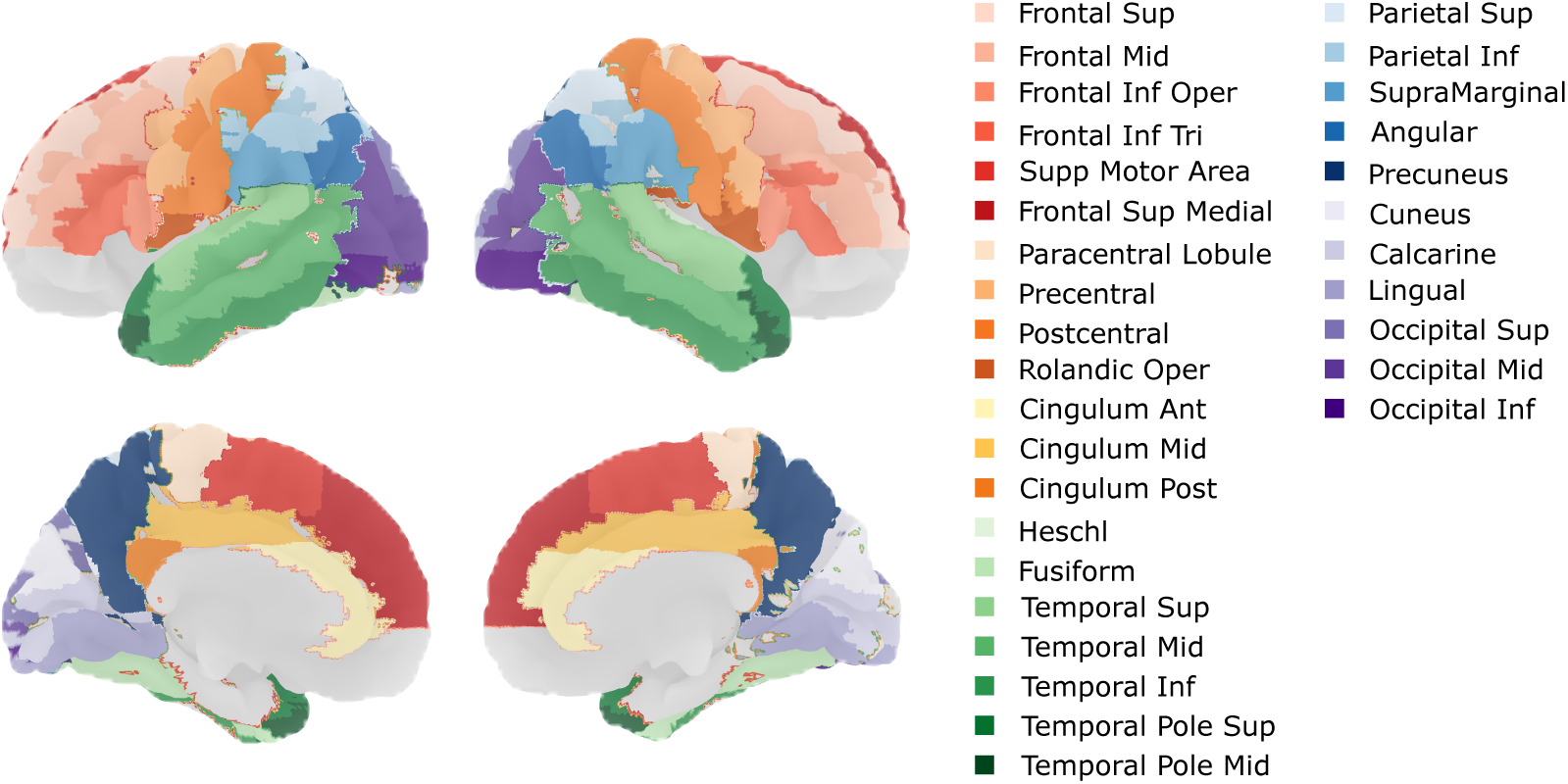
Modified AAL atlas. AAL atlas, from which orbital areas, deep nuclei and cerebellum had been removed, was utilized in the present study.

### 2.4 Power spectra

Power spectral density (PSD) was used for fingerprinting in parallel with the FC metrics because it is a robust metric and has already been shown to distinguish both families and individuals (Leppäaho et al., 2019; Haakana et al., 2024). For PSD calculation the ROI signals were cut to 180 s. Power spectra were calculated by Welch’s method with four-second windows and 50% overlap. From the acquired PSD, average power within twenty-one frequency bins defined in Leppäaho et al. (2019) was calculated to yield 2436 power estimates for each subject. After exclusion of ROIs located in deep subcortical areas, cerebellum, and near the eyes, 1302 power spectral features remained.

### 2.5 Latent-noise Bayesian Reduced Rank Regression

Let N denote number of observations^1^, M number of independent variables, P the number of features, and K the latent rank. LnBRRR takes the form:

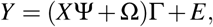

where *Y_N_*_×*P*_ is the observed data matrix that contains flattened FC or PSD data from the subjects, *X_N_*_×*M*_ is the covariate matrix, Ψ*_M_*_×*K*_ is the regression coefficient matrix, Ω*_N_*_×*K*_ is the latent noise matrix, Γ*_K_*_×*P*_ is the mapping from latent space to observation space, and *E_N_*_×*P*_ is the matrix of independent noise (Gillberg et al., 2016).

LnBRRR learns a K-dimensional (K≤M) latent representation of the observed data (*Y*) based on the independent variables. The K latent components learnt by the model are combinations of variables in *Y*, the variation of which the independent variables can explain to a high degree. The weights in row *k* of Γ define how variables in *Y* contribute to the *k*-th latent component. The model allows simultaneous use of multiple independent variables, either categorical or continuous, which sets it apart from simpler models that only utilize a single categorical variable for separating observations, such as LDA. Furthermore, the model enables modeling of weak effects among noise by acknowledging the structured noise in the observed data (Gillberg et al., 2016). This quality makes lnBRRR especially suitable for working with somewhat noisy M/EEG data.

LnBRRR applies shrinkage on Ψ, Ω, and Γ to prevent rotational unidentifiability (Bhattacharya and Dunson, 2011). The columns *k* of Ψ and Ω, and the rows *k* of Γ were given the following priors:

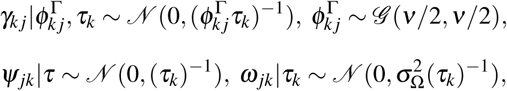

where *N* is the normal distribution and *τ_k_* is defined as:

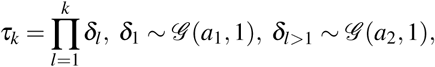

where *G* is the gamma distribution (shape, scale). An extensive description of the lnBRRR algorithm is available in the original publication (Gillberg et al., 2016). The present work utilized a Python implementation (https://github.com/anttoniojanpera/ln-BRRR) of the lnBRRR algorithm which follows the model definition by Gillberg et al. (2016).

### 2.6 Model training

#### 2.6.1 LnBRRR

For the training of lnBRRR, source-level connectomes or PSDs from two recordings per participant were included in the observed data matrix Y. Two observations were used because one observation per participant would fail to convey information on within-participant variation. The Y matrix columns were standardized to zero mean and unit variance. To model individual differences, participants were set as the independent variables. Hence, X contained one-hot encoding of the participants, i.e., each column of X contained binary-coded information on which observations in Y belong to the participant corresponding to that column. Model hyperparameters were selected based on preliminary testing and prior knowledge from previous experiments (Leppäaho et al., 2019; Haakana et al., 2024). The same hyperparameters were used for all metrics, as they yielded acceptable convergence and identification performance. It should be noted that further optimization of hyperparameters would have been possible for each metric, and separately for each frequency band in case of FC metrics, but this would have required substantial additional computational effort and was deemed unnecessary.

Rank 10 and 1000 iterations (50% burn-in iterations) were used for training all the models in the present study. The following hyperparameters^2^ were used in model training: *ν* = 1, *a*_1_ = 2.1, *a*_2_ = 3.1, *a_σ_* = 1000, *b_σ_* = 0.01, and *σ* ^2^ = 1. The Γ matrices were initialized by LDA since LDA has a similar optimization goal to lnBRRR and therefore provides a sensible starting point. Other parameters were initialized with random sampling from the distributions provided above. Gibbs sampler was utilized to sample the posterior distributions of the parameters. Model convergence was established by examination of R-hat values of Ψ, Ω, Γ, local (*ν*) and global (*τ*) shrinkage parameters, and target variable noise parameters. R-hat measures model convergence by estimating the factor by which the dispersion of the posterior distribution might be reduced if iterations were continued *ad infinitum* (Gelman et al., 2013). Median R-hat threshold of 1.1 over four chains, i.e., independent runs of the model with different random initial values (with the exception of Γ), was selected.

#### 2.6.2 PCA and LDA

For lnBRRR benchmarking, PCA and LDA were both trained on resting-state PSD, PLV, iPLV, PLM, and AEC. PCA was trained to find 10 principal components that explain total variation of the observed data instead of individual variation. PCA is theoretically not optimal for fingerprinting as principal components do not separate individual variation from total variation. Furthermore, train-test-splits are not commonly used with PCA as the model does not have a classification/prediction goal. However, PCA was included in the present study because it was of interest whether the principal components also capture individual variation among the total variation. The train-test-splits were utilized to make PCA results comparable to other methods and to assess whether PCA solutions also capture individual variation in unseen data. The mapping between the full data and principal components learnt by PCA was used to project the data to the latent space where cosine distance was used for identification.

LDA was trained to find 10 components that maximally separate the subjects in the training data. To this end, subject identifiers were used as classes. As LDA decision function can only separate unseen observations into the classes involved in training (here, training set individuals), the mapping learnt by LDA was used to project the data to the latent space where cosine distance was used for identification.

### 2.7 Subject Identification

To assess whether lnBRRR successfully captured the individual variation in the training data and to estimate how well the learnt latent solution generalized to unseen individuals, an identification test in the latent space was performed. The FC/PSD data (from two recordings per subject) were mapped to the latent space with generalized inverse of the Γ matrix as was done in previous studies (Leppäaho et al., 2019; Haakana et al., 2024). In the latent space, cosine distances were calculated between the first observation of a subject and second observations of all subjects. Cosine distance was chosen for measuring distance in latent spaces as it is rotationally invariant. The identification was considered successful if the smallest cosine distance was between the first and the second observations from the same subject. In addition, Pearson’s correlation was used as a distance metric alongside cosine distance for the full data because it has been commonly applied in the literature. For Pearson’s correlation, identification was considered successful if the intersubject correlations were smaller than the intrasubject correlation. The identification was performed twice so that the two observations from each individual were both used as the target among other observations, and the mean of the two accuracies was computed. In the present study, identification accuracies were used to determine lnBRRR model performance 1) with low sample sizes, 2) against PCA, LDA and full data, and 3) with task data, as follows:

1. To evaluate whether lnBRRR captures features that generalize beyond the training set with small sample sizes, the lnBRRR identification accuracy was assessed with models trained with N=20-44. Training and test accuracies were computed for each of the two subsets from data split 1, and the mean andstandard error of the mean (SEM) of the resulting two accuracies were reported. The random identification chance was 0.02-0.05 for the training sets, depending on the sample size (N=20-43), and 0.02 for the test set. To further confirm that lnBRRR learnt individual features, lnBRRR accuracies were compared against identification performance in a random space. Random Γ were generated by bootstrap resampling: values of the random Γ row *i* were derived by resampling the values of lnBRRR Γ row *i* with replacement (500 repetitions). This approach guaranteed that random weights have similar distributions as the model weights but lack the assumed individual patterns meaningful for distinguishing individuals. Connectome or PSD data were mapped to random space with the inverted random Γ, and accuracy was computed as with lnBRRR. The null distribution of random accuracies was then used to derive empirical p-values for lnBRRR accuracies as follows: *p* = (*k* + 1)*/*(*n* + 1), where *k* is number of null distribution values equal or larger than the observed lnBRRR accuracy and *n* = 500. Significance threshold of 0.05 was selected, and Benjamini-Hochberg correction was used to address the multiple comparisons problem.
2. To benchmark lnBRRR against PCA and LDA, identification with cosine distance was performed in the latent spaces of each model as described above. Data split 2 was utilized for training all three models to ensure comparability. LDA decision function was additionally used for identification of the training set subjects on PLV data, as cosine distance identification results were unexpectedly low with functional connectivity features. For each CV fold the mean accuracy over the four chains and two identifications were calculated, and the resulting five CV fold accuracies were used to compute the reported mean training and test accuracies and SEM. The random identification chance was 0.02 for the training set and 0.07 for the test set. The differences in test accuracies between the models were statistically assessed by Friedman test so that the models were the classes (treatments) and the accuracies from different metrics were the observations (blocks) (Demšar, 2006). Test statistic (*χ*^2^) and p-value (*p_F_*) were reported. Significance threshold of 0.05 was chosen.

The lnBRRR identification accuracies were also compared to the identification accuracies obtained with the corresponding full connectomes or PSDs. Two approaches for measuring the distance between observations were selected: Pearson’s correlation and cosine distance. The correlation approach enables comparison with identification results from the previous literature whereas the cosine distance enables more appropriate comparison between full data and the latent space accuracies by lnBRRR. Identification was performed as described in the first paragraph of section 2.7. The data was standardized to zero mean and unit variance prior to the identification. Full data accuracies were calculated for training and test sets of the CV folds from data split 2 to enable proper comparison of accuracies with the models. Differences in lnBRRR test accuracies and the full data accuracies on the test set were statistically assessed with Friedman test and with *post hoc* Nemenyi test with significance threshold set at 0.05 and Benjamini-Hochberg correction (Demšar, 2006). Corrected p-values (*p_N_*) were reported for the post hoc Nemenyi test.

1. To assess lnBRRR performance on task data, the model was trained with PSD and PLV data from working-memory, story, and math conditions. Data split 2 was used for training and testing the task models. Model performance was assessed within and between conditions. Within-session accuracy was estimated with unseen data from the same condition and between-condition accuracy with unseen data from a different condition than the condition the model was trained on. For each CV fold the mean accuracy over the four chains and two identifications were calculated, and the resulting five CV fold accuracies were used to compute the reported mean±SEM training and test accuracies. The random identification chance was 0.02 for the training set and 0.07 for the test set. The differences in lnBRRR within-condition test accuracies between the resting-state and task conditions were statistically assessed with Friedman test with significance threshold of 0.05, which corresponds to a critical value *χ*^2^ = 7.800 when the number of treatments (here, task conditions) is four and number of blocks (here, PLV frequency bands and PSD) is seven (Martin et al., 1993). Between-condition test accuracies were also compared with the Friedman test and *post hoc* Nemenyi test.

### 2.8 Comparison of model solutions

In factor analysis, rotation of the acquired factor solutions is common practice. LnBRRR can also be viewed from factor analysis perspective, in which case the Γ matrix corresponds to a factor loading matrix. Therefore, the latent component weights in Γ matrices can also be compared between models with cosine similarity (*S_C_*). For comparison based on cosine similarity, two critical values have been suggested: if *S_C_*≥ 0.95 loadings are practically equal and if *S_C_*≥ 0.85 loadings are similar (Lorenzo-Seva and Ten Berge, 2006). The present study utilized these values as well as a third value (*S_C_* ≥ 0.68 = poorly similar) to evaluate the similarity between models from different training sets and task conditions. Note that the poorly similar components should not be considered similar (Lorenzo-Seva and Ten Berge, 2006) but were included to distinguish dissimilar components with a tendency toward similarity from those with no similarity.

The similarity between latent solutions is not straightforward to measure. To avoid rotational unidentifiability, lnBRRR employs shrinkage to enforce Γ matrix into a configuration that maximizes the variance explained by the first components. This characteristic could cause models trained on different datasets (with e.g., different subjects or conditions) to appear dissimilar only due to a difference in rotation. Rotational differences may be more notable while working with small training sets as the model is more likely to overfit to the training data. Hence, it is essential to align the Γ matrices while evaluating their similarity.

To overcome the rotational differences, Γ were aligned with orthogonal Procrustes analysis. Orthogonal Procrustes analysis finds an orthogonal rotation or rotation-reflection matrix T that best aligns matrices A and B by minimizing sums of squares matrix *E* = *AT* − *B* (Schönemann, 1966). Orthogonal

Procrustes allows only rotation and reflection to align the matrices. Rotation matrix T is unique, if *S*^′^*S*, where *S* = *A*^′^*B*, has distinct eigenvalues (Schönemann, 1966). The uniqueness of T was confirmed for all Procrustes analysis results and only results with unique rotation matrices were reported. Figure 2 presents a schematic example of orthogonal rotation, Procrustes analysis, and alignment of model solutions.

**Figure 2.**
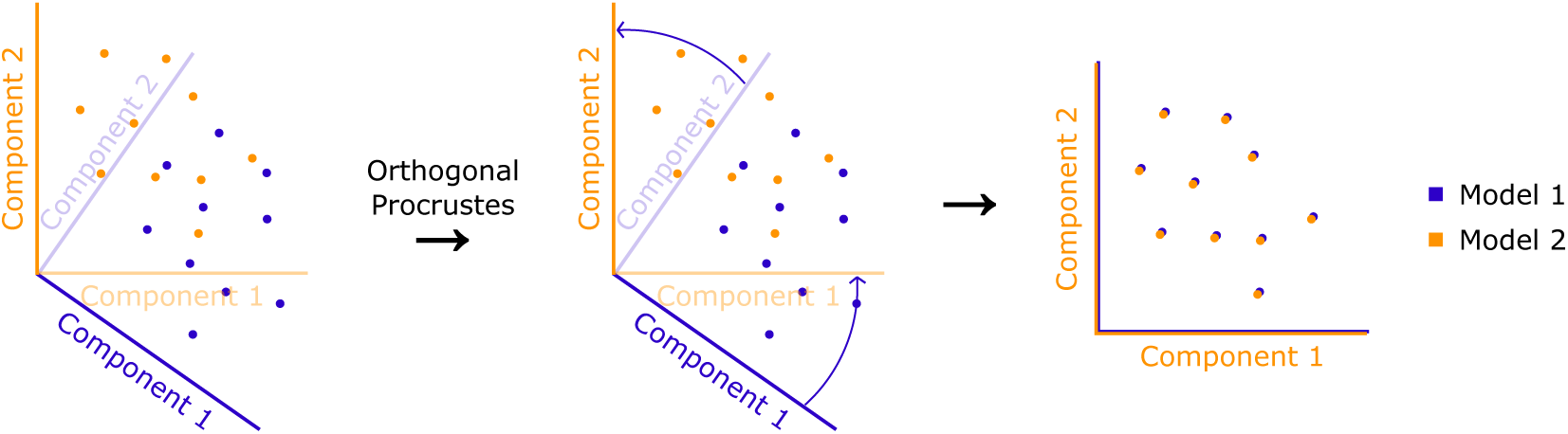
Schematic example of orthogonal rotation. The weights of two components from two arbitrary models are visualized as scatter plots. Orthogonal Procrustes analysis is used to find optimal rotation to align the model weights. Rotational alignment unveiled that the two models were similar up to rotation.

Two comparison scenarios – (1) same task with different individuals and (2) different tasks with same individuals – were examined in the present study. Scenario 1: Models trained on either resting-state PSD or gamma-band PLV from separate training sets were compared. Training sets defined in data split 1 were used. Whereas the identification test on these models reveals lnBRRR generalizability with low sample sizes, the model similarity assessment reveals whether lnBRRR finds reproducible latent individual patterns with low sample sizes. Finding similar patterns regardless of the training set is essential in order to make more reliable neurophysiological interpretations based on the latent components. Scenario 2: LnBRRR was trained on PSD and gamma-band PLV from resting-state and task conditions. Participants with data from all four conditions (N=68) were used for training the model. The comparison of model solutions from different task conditions reveals whether latent individual patterns are affected by the cognitive processes that vary between the tasks.

In both scenarios, pairwise cosine similarities between the corresponding rows of orthonormalized Γ were computed before and after rotational alignment to assess the similarity between the Γ matrices. To ensure that the observed similarities were not found by chance, values in the individual rows of the to-be-rotated Γ were shuffled before Procrustes analysis and the 99.9th quantile of 1001 permutations was selected as the threshold for acceptable post-Procrustes correlation.

### 2.9 Exploratory visualization of latent individual features at brain level

The Γ learnt by lnBRRR is a linear mapping between the latent space and the observed data, and is similar to loading matrices in e.g., PCA or factor analysis. Therefore, it can be interpreted relatively intuitively. In the present study, each latent component can be interpreted as a multivariate pattern of functional connections or power spectral densities that varies between individuals. The component weights in Γ can be visualized at brain level to reveal whether the weights have a meaningful spatial distribution.

To examine power spectral latent individual patterns from different task conditions at the brain level, average weights across the frequency bins were computed for each ROI and those resulting average weights were visualized. Unrotated standardized weights (Z-scores) from the models trained for the similarity testing were used. Note that each component was standardized individually as the shrinkage applied by lnBRRR progressively diminishes the weights of the latent components and makes standardization across components unfeasible.

Considering that only the HCP dataset – a relatively small dataset by machine-learning standards – was used to learn the individual features, the present visualizations remained largely exploratory. To avoid presenting irreproducible latent individual patterns, the number of visualized components was based on the reproducibility results with N=43/44, and only the first three components were visualized. PLV model weights were not visualized as lnBRRR could not find reproducible components for PLV with different training sets.

## 3 RESULTS

### 3.1 LnBRRR retains identification performance with low training set sizes and finds partly reproducible latent power spectral patterns

The first aim in the present study was to evaluate lnBRRR performance with training sets of sizes typical for neuroimaging studies. LnBRRR was trained on PLV (gamma band) and power spectral density with training sets of N=20-44. An increase in training accuracy was observed below N=35 for power spectra and below N=30 for PLV (Fig. 3). PLV test accuracy decreased below N=30 whereas power spectral test accuracy remained stable even at N=20. Training and test accuracies for both metrics plateaued above N=35 (Fig. 3). The test accuracies were compared against a random baseline to determine statistical significance of model generalizability. All test accuracies of both metrics significantly exceeded random baseline. Mean accuracies, SEMs and corrected p-values are shown in SI Table 1. The increased training accuracies and the decreased PLV test accuracy indicated potential overfitting below N=35.

**Figure 3.**
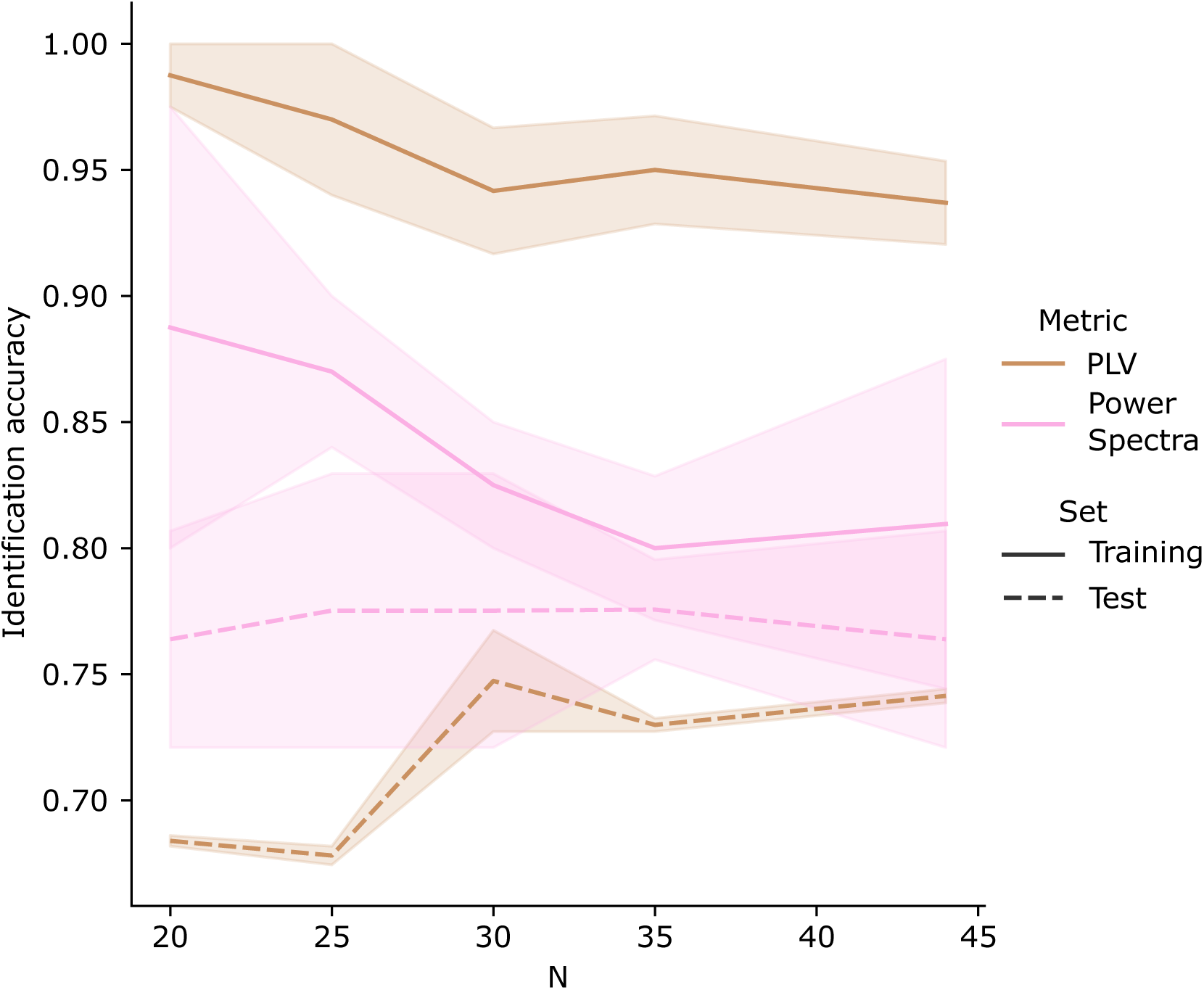
LnBRRR captured generalizable individual power spectral and PLV features with typical neuroimaging sample sizes. Sample size limitations of lnBRRR were assessed with models trained on 20, 25, 30, 35, and 43/44 subjects. PLV in gamma frequency band and power spectra were used for training. Mean identification accuracies and SEM over two training and test sets are shown. Test accuracies were tested against random accuracy baseline. Empirical p-values corrected by Benjamini-Hochberg method were computed for the test accuracies to evaluate model generalizability. All test accuracies for both metrics reached significant (p*<*0.05) corrected p-values. Random chance was ∼0.05-0.02 (N=20-43/44) for the training set and ∼0.02 for the test set.

**Table 1.**
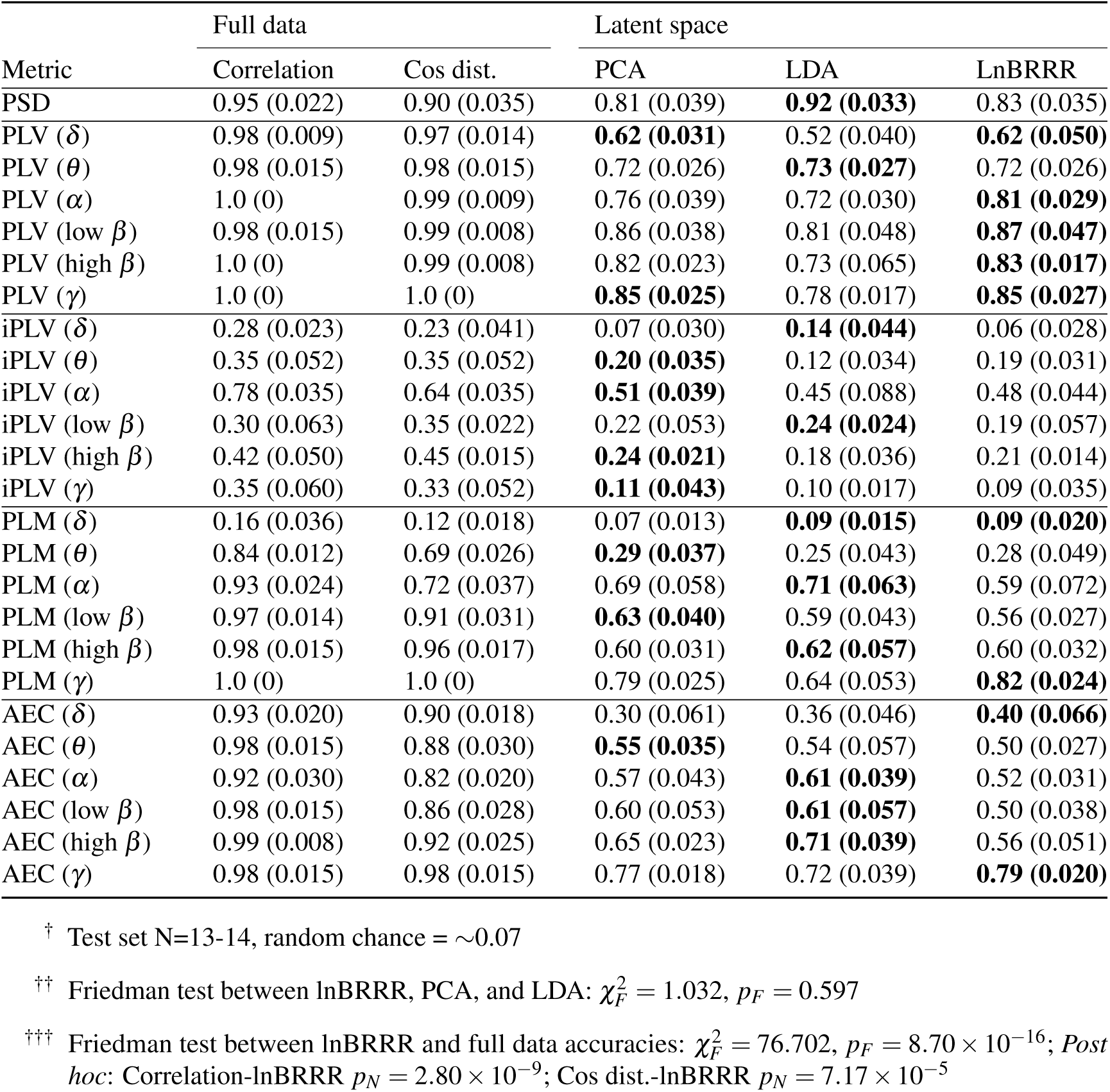
Mean accuracies (SEM) on the test set with full data and latent spaces. Full data identification was performed with Pearson’s correlation (Correlation) and cosine distance (Cos dist.) as the distance function. In the PCA, LDA, and lnBRRR latent spaces, the identification was performed with cosine distance as the distance function.

The similarity between resting-state lnBRRR solutions trained with separate training sets was tested with N=20-44 to establish how reproducible the individual latent patterns found by lnBRRR are. Initially, power spectral models were dissimilar with all training set sizes (Fig. 4). The rotation by Procrustes analysis revealed that the number of similar components for the power spectral models was one with N=20-25, two-three with N=30-35, and three-four with N=43-44 (Fig. 4). Several components also showed tendency toward similarity at all training set sizes (Fig. 4). PLV models derived from two separate training sets were not similar before or after rotation with the largest training set (SI Fig. 1). This indicates that latent solutions for individual differences in power spectral density get increasingly reproducible as the sample size is increased whereas for PLV consistent solutions are substantially harder to acquire.

**Figure 4.**
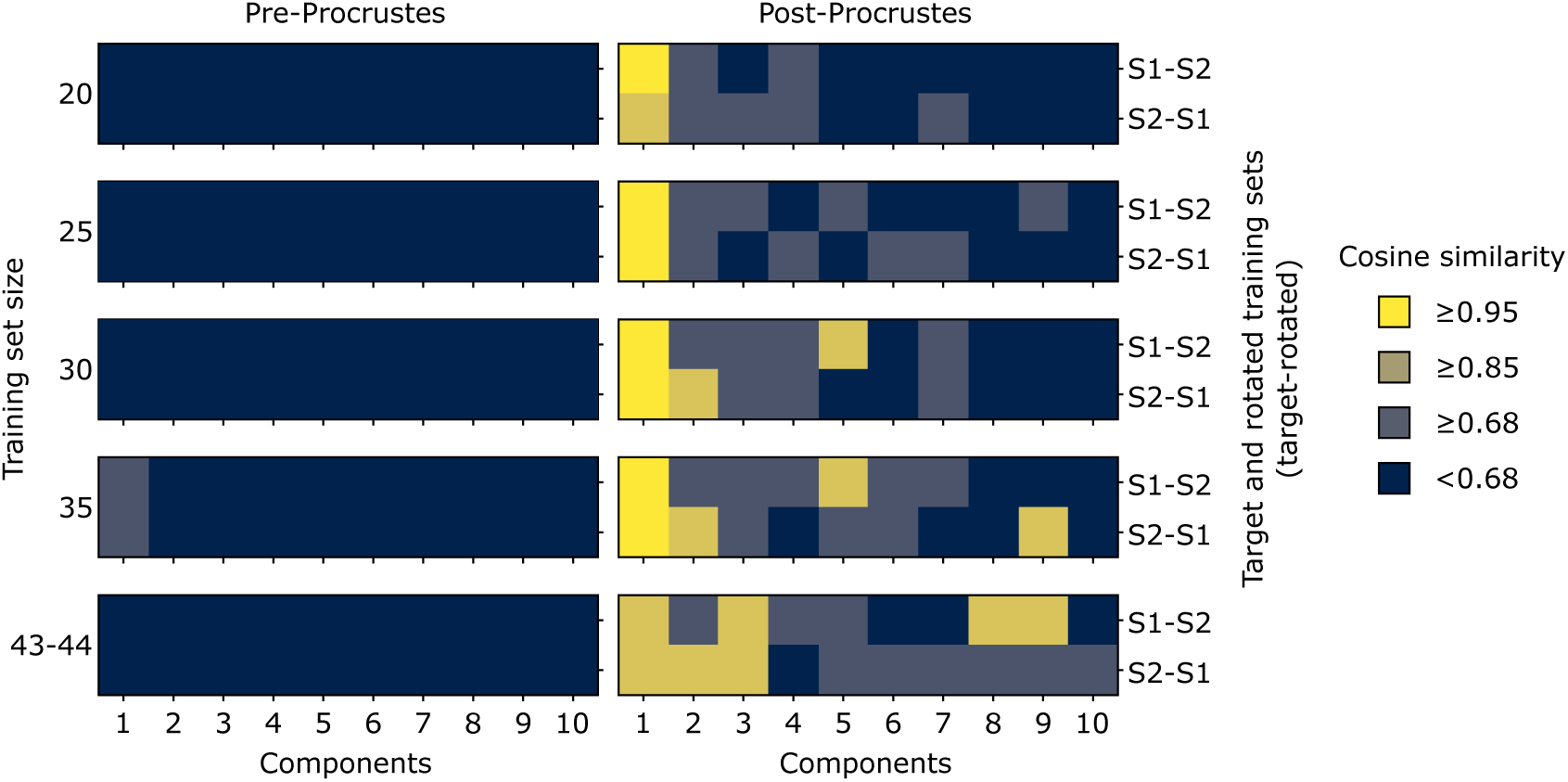
The reproducibility of resting-state power spectral lnBRRR solutions increased with training set size. LnBRRR reproducibility w.r.t. training set size was assessed by measuring the similarity between model solutions acquired with different training sets (N=20-44). Similarity was measured by cosine similarity between orthonormalized Γ matrix rows before and after rotational alignment by Orthogonal Procrustes analysis. In each individual heatmap, x-axis indicates the Γ rows (Components) and y-axis indicates the target and rotated training sets (S1=set 1, S2=set 2) examined. Cosine similarity was divided into four classes: practically equal (≥ 0.95), similar (≥ 0.85), poorly similar (≥ 0.68), and no similarity (*<* 0.68).

### 3.2 LnBRRR performs comparably to PCA and LDA

To benchmark lnBRRR against PCA and LDA the models were trained on PSD and four FC metrics (PLV, iPLV, AEC, PLM) and subject identification was used to assess how well each model captured individual features. LnBRRR attained overall higher test accuracies over the frequency bands with PLV than with the other FC metrics (Table 1). PLV test accuracies were approximately 0.8 on all but delta and theta frequency bands (Alpha: 0.81±0.029, Beta low: 0.87±0.047, Beta high: 0.83±0.017, Gamma: 0.85±0.027), whereas AEC and PLM reached comparable accuracies only in the gamma band (AEC: 0.79±0.020, PLM: 0.82±0.024) (Table 1). Imaginary PLV performed poorly on all frequency bands. Power spectral model performed comparably to PLV models reaching test accuracy of 0.83±0.035.

Comparison of lnBRRR performance to PCA and LDA revealed no significant differences between the models on the test data (*χ*^2^ = 1.032, *p* = 0.597). The corresponding training accuracies for all methods and metrics are listed in the supplementary information (SI Table 2). LDA training accuracies with functional connectivity were ill-behaved as low training accuracies but high test accuracies were observed (Table 1, SI Table 2). Similar behavior was not observed with power spectral LDA. LDA is optimized with Mahalanobis distance, so it may behave unexpectedly with other distance functions. To test whether difference in distance function could be the source of the ill-behaved performance, LDA decision function was used to identify the training set individuals on PLV data. LDA decision function achieved very high training accuracies and surpassed cosine distance accuracies with all PLV frequency bands (SI Table 3). This demonstrated that cosine distance may be poorly suited for LDA latent spaces in some cases.

To compare lnBRRR performance to the accuracies acquired with full functional connectomes and PSD, full data accuracies were calculated for the FC metrics and PSD without dimensional reduction. Although lnBRRR reached test accuracies relatively close to those of full data with power spectral density and the best performing frequency bands of PLV, PLM, and AEC (Table 1), the lnBRRR test accuracies remained significantly lower than the full data accuracies with the test data (*χ*^2^ = 76.702, *p_F_* = 8.70 × 10^−16^; *Post hoc*: Pearson correlation-lnBRRR: *p_N_* = 2.80 × 10^−9^; Cosine distance-lnBRRR: *p_N_* = 7.17 × 10^−5^). Full data test accuracies with Pearson’s correlation and cosine distance did not differ significantly in the *post hoc* analysis (*p_N_*= 0.117).

### 3.3 LnBRRR solutions from task data generalize to unseen individuals and across task conditions

To assess the extent to which lnBRRR captures individual differences from task data, the model was trained with PLV and PSD from resting-state, working-memory (WM), story and math tasks. The task affected which frequency bands generalized better to unseen individuals (Fig. 5). High-beta and gamma bands were better for working-memory task (High-beta: 0.92±0.031, Gamma: 0.91±0.028) whereas for story and math tasks the alpha band was somewhat pronounced (Story: 0.87±0.024, Math: 0.81±0.084) (Fig. 5). Power spectra performed comparably to the best generalizing PLV frequency band on each task (WM: 0.93±0.017, Story: 0.86±0.048, Math: 0.77±0.041 (Fig. 5). There was no significant difference in lnBRRR performance between the tasks (*χ*^2^ *_F_*= 5.870, critical *χ*^2^ *_F_*= 7.800).

**Figure 5.**
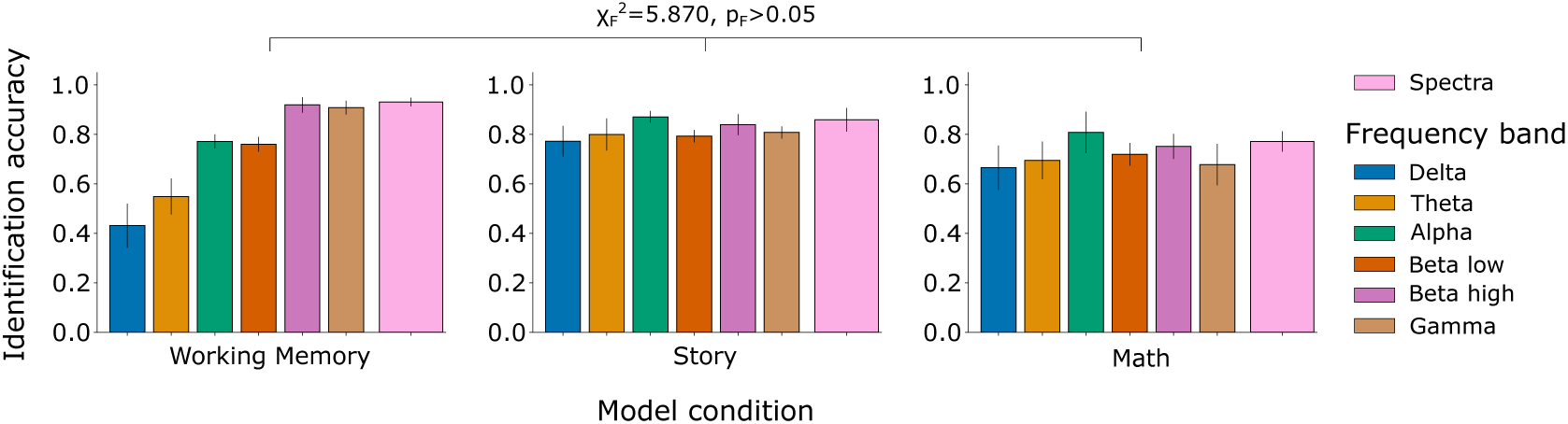
LnBRRR performance remained on a comparable level across the task conditions. LnBRRR was trained with PLV and power spectra data from working-memory, story, and math tasks. Mean test identification accuracies and SEM over the CV-folds are shown. Random chance was ∼0.07 for the test set. LnBRRR performance between resting-state (not shown) and task conditions was tested with Friedman test. LnBRRR performance did not differ significantly between the resting-state and task conditions (*χ*^2^ = 5.870, *p_F_ >* 0.05).

Model generalizability across task conditions was examined by identification of individuals based on their resting-state PLV and power spectral data with lnBRRR models trained on a different task conditions (WM, Story, Math). The obtained cross-condition accuracies were high overall (Fig. 6), and no significant difference in test accuracy was observed between the resting-state and task models while identifying individuals with resting-state data (*χ*^2^*_F_* = 7.922, critical *χ*^2^ *_F_*= 7.800; *Post hoc*: Rest-WM: *p_N_* = 0.586; Rest-Story: *p_N_* = 0.586; Rest-Math: *p_N_* = 0.285; WM-Story: *p_N_* = 1.000; WM-Math: *p_N_* = 1.000; Story-Math: *p_N_* = 1.000). Cross-condition identification was also tested with task data, and the results were overall similar to the results presented here with resting-state data (SI Fig. 2). The high cross-condition accuracies imply that weighting matrices (Γ) learnt by lnBRRR are relatively similar regardless of the task condition.

**Figure 6.**
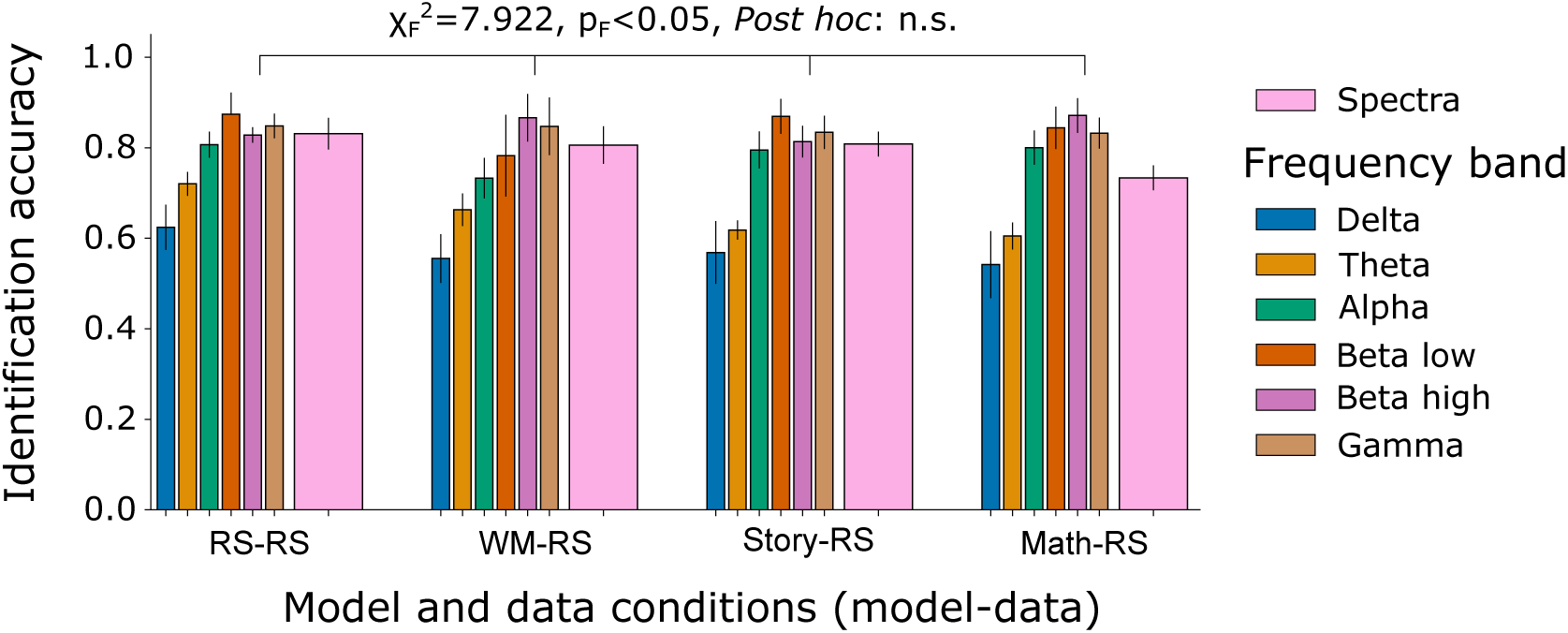
Task lnBRRR models attained comparable identification performance to the resting-state model with resting-state data. Resting-state PLV and power spectra data, and lnBRRR models trained on data from each condition were used to assess whether lnBRRR models trained on task data can identify individuals based on their resting-state data. Mean test identification accuracies and SEM over the CV-folds are shown. The task condition of the model used for identification is shown on the x-axis. Resting-state model (RS) results are shown for comparison. Random chance was ∼0.02 for the training set and ∼0.07 for the test set. LnBRRR performance with resting-state data between the resting-state and task models was tested with Friedman test. LnBRRR performance on resting-state data did not differ significantly between the models from resting-state and task conditions after correcting the *post hoc* p-values (*χ*^2^ = 7.922, *p_F_ <* 0.05, *Post hoc*: n.s.). n.s.=not significant

### 3.4 Rotational alignment reveals high between-task similarity of power spectral lnBRRR solutions

To elucidate the extent of similarity between model solutions from different task conditions cosine similarity was computed for the corresponding rows of Γ from separate models before and after rotational alignment by orthogonal Procrustes analysis. Before Procrustes analysis, Γ of power spectral models from different conditions were not similar as all cosine similarity values were below the 0.85 threshold (Fig. 7). PLV models from story and math conditions had one similar component before Procrustes analysis (SI Fig. 3). The similarity of Γ rows improved notably with rotational alignment for the power spectral models (Fig. 7). Procrustes analysis on power spectral models revealed high similarity between resting-state and working memory conditions as only two non-similar components were found for the pair after Procrustes analysis (Fig. 7). Story and math conditions were also highly similar (Fig. 7). The less similar condition pairs (e.g., working memory vs. story) all had at least five similar components (Fig. 7). Although successful with power spectral models, Procrustes analysis did not markedly improve similarity between PLV models, as only one or two similar components were observed for some condition pairs after Procrustes analysis (SI Fig. 3).

**Figure 7.**
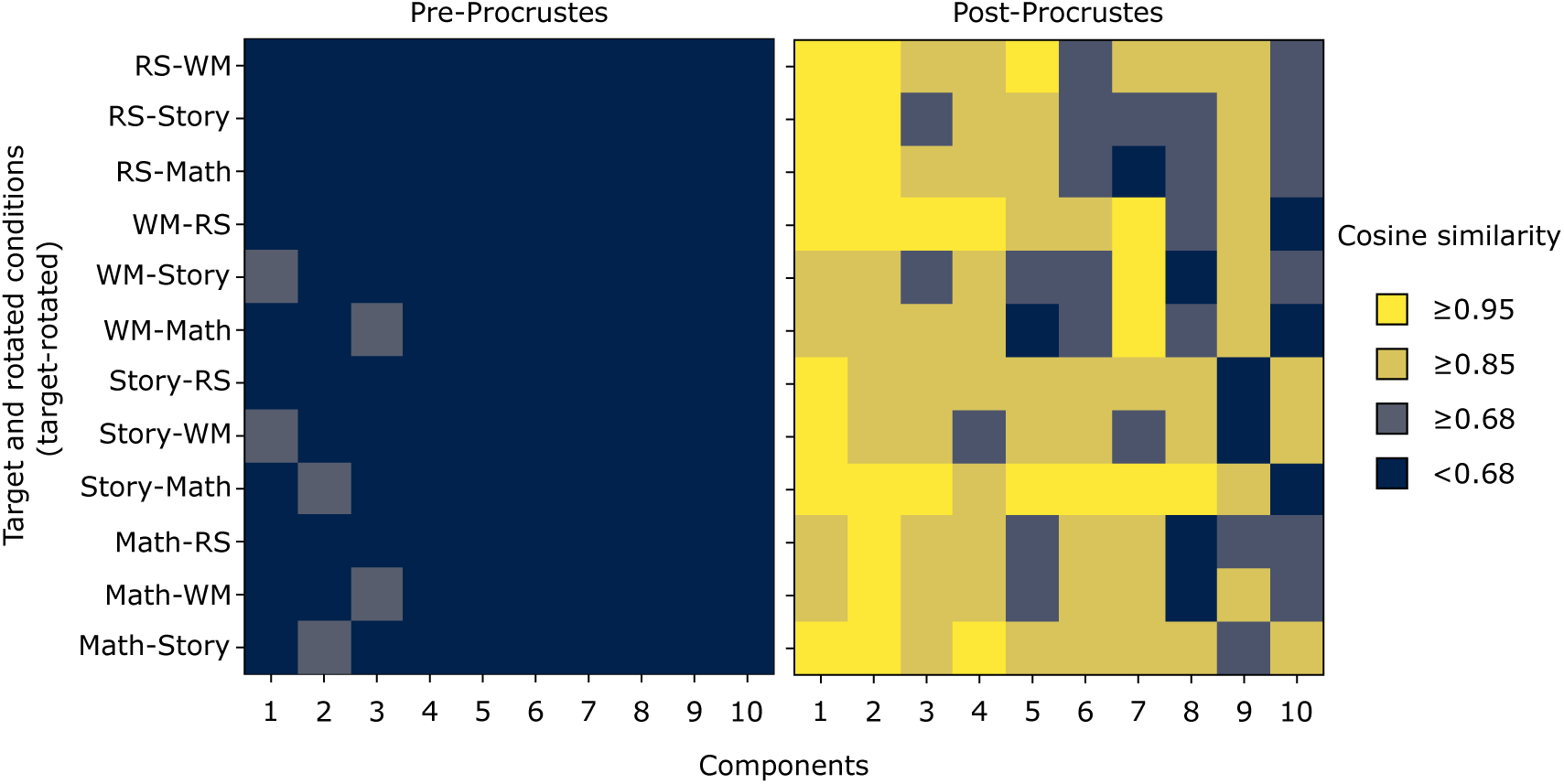
Power spectral lnBRRR models captured largely similar but differently rotated individual features across task conditions. The similarity between lnBRRR solutions from different task conditions was measured by cosine similarity between orthonormalized Γ matrix rows. Similarity was measured before and after rotational alignment by Orthogonal Procrustes analysis. X-axis indicates the Γ rows (Components) and y-axis indicates the condition pair examined. Cosine similarity was divided into four classes: practically equal (≥ 0.95), similar (≥ 0.85), poorly similar (≥ 0.68), and no similarity (*<* 0.68).

### 3.5 LnBRRR weights can reveal spatiospectral patterns that distinguish individuals

Finally, an exploratory visualization of the weights learnt by lnBRRR at the brain level was conducted. Only the first three components of power spectral models from each task condition were shown because three reproducible components were observed when models trained with separate training sets were examined (Fig. 4). PLV model weights were not visualized as no reproducible components were found with N=43-44 (SI Fig. 1). Unrotated weights were visualized to reveal whether solutions from different task conditions showed similarities also without rotational alignment, as was suggested by the high cross-condition identification accuracies. The weights were almost invariably ROI specific rather than frequency bin specific (SI Fig. 4). Because of this, the averaged weights per ROI were visualized. Power spectral model components showed similar weights on both hemispheres with minor hemisphere-specific details apparent (Fig. 8). Component 1 contrasted frontal, and to some extent temporal, areas with parietal and occipital areas regardless of the condition (Fig. 8). Component 2 was more variable between conditions, but resembled Component 1 with minor differences (Fig. 8); this was confirmed by comparison of all weights (SI Fig. 4). Component 3, apart from the story task, weighted predominantly the cingulum (Fig. 8). Altogether, the results demonstrated that latent individual power spectral features from different conditions appeared relatively similar without rotation, but closer examination revealed that individual weights varied more notable between conditions. All models contrasted frontal and temporal regions with parietal and occipital regions, in addition to which cingulum was almost invariably pronounced across conditions.

**Figure 8.**
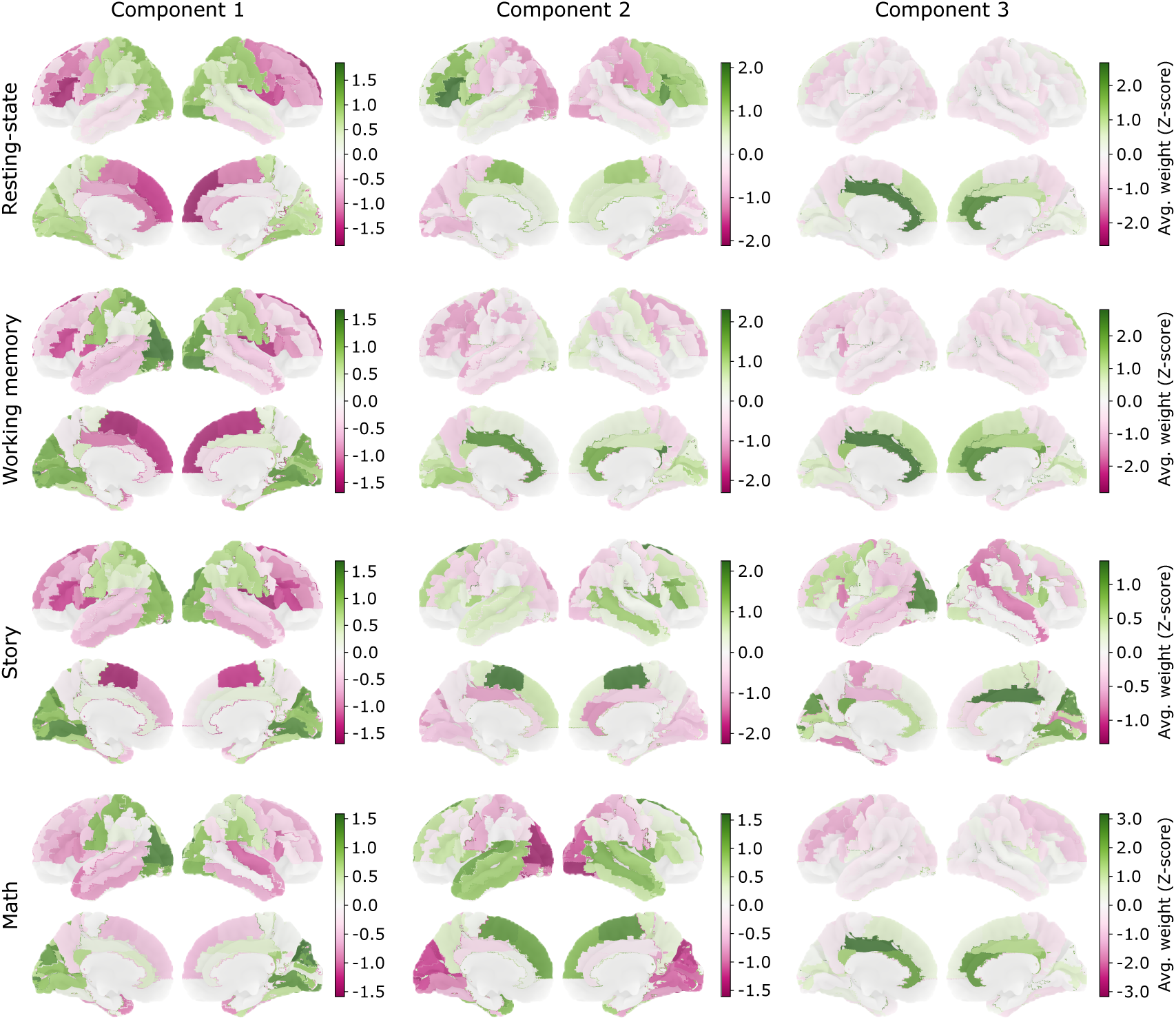
Unrotated average lnBRRR weight patterns were partly similar between the task conditions. LnBRRR was trained on resting-state, working-memory, story, and math task power spectra from the same 68 participants. The average weight over the frequency bins for each ROI of the first three unrotated components are shown for each model. Note that lnBRRR progressively diminishes component weights; thus, the components were scaled individually and the colorbars have different scales between components and conditions.

## 4 DISCUSSION

Direct correlation-based comparison of individual functional connectomes or power spectral densities is fast and distinguishes individuals. Yet, that approach lacks the beneficial qualities offered by modeling approaches. Modeling provides the opportunity to study effects of specific independent variables in the data and can also provide simultaneous dimension reduction. With linear models, the model weights can be intuitively interpreted at the brain level, thus offering control over the effects of interest and a neuroscientifically meaningful solution.

The present work aimed to advance the modeling of individual brain features and addressed lnBRRR fingerprinting performance with low sample sizes and task data while regarding participants as independent variables. Overall good identification performance was observed with resting-state and task data, and accuracy remained high when subjects were identified between task conditions. Sample size reduction towards N=20 caused minor signs of overfitting while N=30-35 was sufficient to retain training and test performances that could not be improved with further increase of the training sample size. Power spectral lnBRRR solutions acquired with low sample sizes showed reproducible latent patterns and lnBRRR solutions trained with power spectral density data from different task conditions could be rotated to a similar configuration, suggesting that one underlying latent solution could explain individual differences in all conditions. In brief, the results demonstrated that lnBRRR is a potential research tool for neuroscience research and can be leveraged even with small sample sizes while utilizing PSD data, and that latent patterns of individual differences in power spectral density remain unchanged by task conditions.

### 4.1 Power spectral latent fingerprints capture generalizable and partly reproducible individual patterns with typical neuroimaging sample sizes

The first aim in the present study was to assess whether lnBRRR could be utilized with sample sizes typical for neuroimaging literature. LnBRRR was trained with N=20-44, and identification accuracies were computed with training and test sets to assess signs of overfitting. Training accuracies were increased when sample size decreased below N=35 and N=30 for power spectra and PLV, respectively. These increases could be inflated due to increased random chance as the training set size decreased. However, the random chance increased by approximately 3 percentage points when sample size was decreased from 44 to 20 while the training accuracies of PLV and PSD simultaneously increased by 5 and 8 percentage points, respectively. Thus, overfitting is likely to, at least partially, cause the increased training accuracies. PLV test accuracy started to decrease below N=30 whereas power spectral test accuracy remained stable regardless of the training sample size. All test accuracies from both metrics significantly exceeded the random baseline which demonstrates lnBRRR to be functional even at N=20. Nevertheless, training sample size of 30-35 or higher is recommended to prevent possible overfitting and to obtain optimal performance. It is also noteworthy that low sample size may exacerbate convergence issues while training; hence, gathering as large a sample as possible is recommended. These results demonstrate that although the choice of metric can slightly affect model performance with low sample sizes, lnBRRR remains overall functional at sample sizes typical for neuroimaging experiments.

In addition to generalizability, the reproducibility of the latent solutions was examined. Similarity testing on resting-state data from different training sets revealed that power spectral models had one similar component at N=20 and the number increased to three-four at N=43-44. Several components tended toward similarity suggesting that with increased sample size more reproducible components could be extracted. The reproducible components together with generalizable identification performance that exceeded random baseline imply that the power spectral individual patterns found by the model are neurophysiologically meaningful.

PLV models remained dissimilar regardless of rotational alignment even at N=43/44. The dissimilarity between PLV lnBRRR solutions contradicts the excellent identification performance in unseen individuals. The reproducibility difference between PSD and PLV is likely explained by the difference in the complexity of the metrics. Power spectral density is a simple and robust metric that gives a relatively broad summary of brain activity across the brain whereas PLV gives a much more detailed description of interareal synchronized activity. Due to the enormous number of possible connections and networks, it is feasible that individual differences in the functional connectivity networks can be captured into several dissimilar latent solutions that reflect individual differences in separate networks but still generalize to unseen data. The successful comparison against the random baseline further implies that the captured latent PLV features are, at least in part, neurophysiologically meaningful, albeit not reproducible. Moreover, the low sample size could exacerbate the reproducibility problems, as the high number of features (functional connections) and low training set size likely cause the model to favor individual differences that are prominent in the training data but may be poorly generalizable. Furthermore, it has been shown that factor analysis with six factors requires a sample size of 75-110, depending on communality and variables to factors ratio, and that increasing variables to factors ratio beyond six does not decrease the required sample size (Mundfrom et al., 2005). Therefore, considering the rank (10) used in the present study and lnBRRR model complexity, a larger sample size would likely be required to reliably discover generalizable and reproducible distinguishing factors between individuals. A subset (N=68) of HCP dataset was selected for the present study because HCP MEG data has already been examined in fingerprinting context and because HCP data contains a notably large sample of subjects with MEG recordings from several task conditions. Future efforts with larger datasets are required to further unveil individual functional connectivity features from brain data.

### 4.2 LnBRRR reaches comparable accuracies to alternative dimension reduction methods

As the next step, lnBRRR performance was compared to PCA and LDA. Statistical testing showed that lnBRRR latent space captured individual features comparably to PCA and LDA in terms of identification accuracy. These results imply that simpler methods suffice for neural fingerprinting considering that PCA and LDA are computationally lighter and can be interpreted based on the mappings from data to latent space. In more complex research settings, however, lnBRRR is a strong candidate for fingerprinting due to its beneficial characteristics. LnBRRR enables more diverse conditioning of the latent space through multiple independent variables and finds small effects in noisy data, which supports its use in individualized neuroimaging and diagnostics over PCA or LDA (Gillberg et al., 2016). Although not put to test in the present study, lnBRRR has previously been demonstrated to work with multiple independent variables (Gillberg et al., 2016; Leppäaho et al., 2019). Future research efforts should experiment more in depth the capabilities of lnBRRR with multiple independent variables in the neuroimaging setting, e.g., by distinguishing between individuals and medical condition groups simultaneously. It is also possible that identification accuracy in the lnBRRR latent space could be further increased by additional optimization of the model hyperparameters or the classification method used for identification, e.g., by using support vector machines or other supervised machine learning models, yet such experiments were out of the scope of the present study.

Cosine distance was observed to be a suboptimal metric for identification in LDA latent spaces, as the functional connectivity training accuracies were poor with cosine distance but not with LDA decision function. The problem was likely related to the distance function as LDA decision function performed well for the training set. Since LDA decision function can only be applied for the training set, its use for neural fingerprinting is contradicted, unless future work should identify other distance metrics that are better suited for this purpose.

LnBRRR was also compared against accuracies obtained with full data. The full data accuracies were in line with previous studies (Sareen et al., 2021; Da Silva Castanheira et al., 2021). With 10 latent components, lnBRRR reached high test accuracies but did not quite reach the performance of the full data. LnBRRR training accuracies were closer to the full data accuracies.

LnBRRR performance was comparable to the previously published results by Haakana et al. (2024). As before, PLV achieved overall the best performance, and gamma-band PLM and AEC also reached fair accuracies whereas iPLV identification performance was poor overall. Power spectral density proved again to be a robust metric for fingerprinting (Haakana et al., 2024). Notably, in the present study the lnBRRR model with 10 latent components achieved nearly the same level of performance as was previously obtained with 20 components (Haakana et al., 2024). It is possible that part of this improvement was due to model training differences, as Haakana et al. (2024) used only one observation per subject for training whereas two observations were utilized in the present study. Furthermore, the relatively high random chance (∼0.07), due to the small CV folds, could have slightly inflated the test accuracies.

### 4.3 Latent fingerprints derived from power spectral data capture largely similar individual patterns across task conditions

No statistical difference in lnBRRR performance was observed between different task conditions. The task condition affected which frequency bands of PLV generalized better to unseen individuals. For the working-memory task, the lower frequency bands had poorer test accuracies while high beta and gamma band test accuracies showed better generalization. Alpha band test accuracy was pronounced in the story and math conditions. Power spectral models performed well in all task conditions and resulted in comparable accuracies to the resting-state condition. The results imply that task condition does not affect the identification performance. MEG data from task conditions with a strictly controlled procedure (e.g., mismatch negativity, auditory steady state responses) have previously been shown to provide better identification results compared to more leniently controlled tasks (e.g., prose listening) (Colenbier et al., 2023). In the present study, the story and math tasks both contained a longish listening period (3°0 s) and could be classified as a leniently controlled task, whereas working-memory task could be considered a strictly controlled procedure. The present findings did not agree with Colenbier et al. (2023), as no difference in performance was found across the tasks. Colenbier et al. (2023) also observed resting-state accuracies to be poorer than accuracies of any task condition, which is also discrepant with the present findings. Colenbier et al. (2023) utilized two separate resting-state recordings and one task recording split into two parts in their testing, leading to possible inflation of task accuracies. This could explain the differences with the results of the present study.

Cross-condition identification was furthermore used to assess the lnBRRR generalizability across conditions. All task lnBRRR solutions for PLV on alpha, low-beta, high-beta, and gamma bands as well as for PSD reached high accuracies when resting-state data was used in identification, whereas cross-condition performance was more modest with PLV on delta and theta bands. Friedman test between the resting-state and task models confirmed that all models performed comparably with the resting-state data. In addition, all models were shown to successfully identify individuals regardless of the data condition, nearly reaching the performance of the model trained on the data condition. Cross-condition identification has been examined in previous research: Finn et al. (2015) showed that identification across conditions with fMRI FC data can reach up to 0.93 accuracy and Gratton et al. (2018) observed that fMRI FC networks from the same and different conditions are similar within individuals. Colenbier et al. (2023) observed high accuracies between rest and several task conditions with PLM and power spectra derived from MEG data. These observations agree well with our present findings using lnBRRR modeling.

LnBRRR generalizability across task conditions suggested that the individual latent patterns captured by the model could be relatively similar across the task conditions. Haakana et al. (2024) quantified similarity of lnBRRR solutions by computing correlations between the latent spaces with Mantel test. This method, however, induces ambiguity to the similarity assessment as both the pseudo-inverted Γ matrix and the observed data affect the latent solution. As Γ can be interpreted as a factor loading matrix in factor analysis, we chose a more direct approach to model comparison by computing cosine similarity between the rows of Γ matrices to assess the similarity of component loadings. Additionally, Procrustes analysis was utilized to assess whether the solutions are similar but differently rotated. Initially, the components were dissimilar with isolated similar components observed between PLV models. Rotational alignment revealed that power spectral lnBRRR solutions shared 5-9 similar components whereas PLV solutions remained dissimilar even after rotational alignment. The results imply that spatiospectral individual patterns remain relatively stable regardless of the task condition, as the task conditions mostly affected the rotation of the latent components. Congruent with the present results, Gratton et al. (2018) found that the similarity between fMRI connectomes derived from multiple recordings and conditions was better explained by the individual than the task condition, suggesting that fMRI functional connectivity networks remain largely similar within individuals even during tasks. Altogether, the present and previous findings align with the view that brain function is mostly intrinsic, and task conditions induce increases or decreases to this activity at specific regions (Raichle, 2010). Therefore, the similar spatiospectral patterns found by lnBRRR across task conditions have plausible neurophysiological basis.

As mentioned above, the apparent lack of similarity of PLV solutions could be caused by the high complexity of brain functional connectivity networks, which could not be sufficiently captured to 10 components, or which would need more data to be captured appropriately. Evidently, it is also possible that the observed dissimilarity is plainly caused by different networks being active during different tasks. Future studies should aim to disentangle whether the dissimilarity between latent solutions is caused by strong task-dependent differences in PLV-derived networks or merely insufficient sample sizes for finding reliable latent representations. This would likely require large datasets (N*>>*100) with multiple task conditions.

### 4.4 LnBRRR leverages differences between broad areas to distinguish individuals

The average power spectral lnBRRR weights for the first three components were examined for each condition. The weights were not specific to any frequency bin; instead, mostly similar weights were given to all frequency bins within the ROIs. The power spectral model components weighted ROIs within broad brain regions, e.g., temporal lobe, with similar weights. The models contrasted frontal and temporal regions with parietal and occipital regions to some extent in all conditions, and highlighted cingulum in all conditions except the story task. The juxtaposition of frontal and temporal areas with parietal and occipital areas implies that individuals could exhibit an inter-individually variable anterior-posterior balance in the power between these areas, and that lnBRRR can leverage this balance to distinguish individuals. Similar anterior-posterior juxtaposition in lnBRRR weights was found in previous work where familial power spectral differences were examined (Leppäaho et al., 2019). Somewhat congruent anterior-posterior power spectral distributions in resting-state MEG have been found in the peak frequencies of theta, alpha, and beta bands as well as between transient states extracted with Hidden Markov Modeling (Vidaurre et al., 2018; Mahjoory et al., 2020; Guichet et al., 2025). In theory, individual differences in the peak frequency gradients or durations of the transient states could, at least in part, explain the observed spatial distribution of lnBRRR weights; however, this remains to be confirmed.

The components that put weight on the cingulum gave minimal average weights to other areas. This weighting behaviour is in line with previous findings that showed cingulum power in frequency bands from delta to gamma to be salient within individuals, whereas the power spectral saliency was more varied over the frequency bands (Da Silva Castanheira et al., 2021). Functional connectivity results from previous studies demonstrated that cingulum contributes to correlation-based identification of individuals in the gamma frequency band of PLV and AEC (Colenbier et al., 2023; Sareen et al., 2021), and that middle and posterior cingulum, as well as frontal cingulum, depending on the task, were involved in identification with task data (Colenbier et al., 2023). Cingulum has been associated with various highly individualized functions such as executive control and emotion (Bubb et al., 2018), which could explain its importance for distinguishing individuals.

Direct comparison between present work and previous reports is complicated by methodological differences, as lnBRRR does not highlight within-individual saliency but creates combinations of the observed data (i.e., latent components) that are explained by the independent variables. Nonetheless, the consistent observation on the importance of cingulum for distinguishing individuals demonstrates that lnBRRR latent spaces may capture the same neurophysiological phenomena as the intraclass correlation used in previous studies.

### 4.5 Limitations and future prospects

The present findings elucidate the limitations and capabilities of lnBRRR modeling on neurophysiological data and provide a novel, direct method for comparison of linear projection methods in the neural fingerprinting context. Still, several questions remain for future studies to address. LnBRRR has been tested with a limited number of datasets. LnBRRR has been shown by the present and previous studies to work well on datasets that are typical or relatively large in size from the human neuroimaging perspective but would be considered very small from the machine learning point of view (Leppäaho et al., 2019; Haakana et al., 2024). Recent efforts have also demonstrated that lnBRRR can handle large neuroimaging datasets and find latent components that distinguish individuals better than the full data (Heikkinen et al., 2026). Hence, future efforts should focus on putting the model to further test with datasets consisting of hundreds of individuals. The possibility to generalize results to unseen data provided by the modeling approach should be tested among multiple large data sets. Such an inquiry would provide more reliable and thorough understanding of the connections or power spectral values most variable between individuals. Additionally, the stability of fingerprints over longer time periods should be examined since a large bulk of neural fingerprinting literature has utilized recordings from one session or even a single recording divided into multiple parts. Furthermore, lnBRRR should be put to test in tasks beyond individual identification to estimate its potential for use in diagnostics. Classification of participant groups based on age or health status or predicting behavioral variables are examples of such tasks.

## CONCLUSION

To summarize, the present study showed that lnBRRR can learn low-dimensional latent spaces that distinguish individuals based on MEG FC and power spectral metrics. The model successfully created generalizable latent spaces with low sample sizes (N=20-44) and learnt partly reproducible latent individual patterns with power spectral density. Viability under small sample sizes makes lnBRRR a powerful tool for neuroimaging data analysis, yet the results obtained with low sample sizes should not be too eagerly generalized beyond the training sample. Latent spaces derived from task data distinguished individuals comparably to resting-state latent spaces and lnBRRR latent solutions generalized between the task conditions. Power spectral latent individual patterns were shown to be largely similar across task conditions. This finding elucidated why lnBRRR solutions generalize well between conditions and provided support for the notion that lnBRRR captures neurophysiological phenomena that vary most between individuals. Altogether, the present results suggest that individual power spectral features are largely intrinsic and remain unchanged by varying cognitive processes.

## AUTHOR CONTRIBUTIONS

**Joonas Karhula**: Writing – original draft, Writing – review & editing, Formal analysis, Visualization, Conceptualization, Methodology. **Anttoni Ojanperä**: Writing – review & editing, Software, Methodology. **Ersin Yılmaz**: Writing – review & editing, Software, Methodology. **Susanne Merz**: Writing – review & editing, Conceptualization, Methodology. **Samuel Kaski**: Writing – review & editing, Methodology. **Riitta Salmelin**: Writing – review & editing, Supervision, Conceptualization, Methodology.

## ACKNOWLEDGMENTS

Data were provided by the Human Connectome Project, WU-Minn Consortium (Principal Investigators: David Van Essen and Kamil Ugurbil; 1U54MH091657) funded by the 16 NIH Institutes and Centers that support the NIH Blueprint for Neuroscience Research; and by the McDonnell Center for Systems Neuroscience at Washington University.

The calculations presented above were performed using computer resources within the Aalto University School of Science “Science-IT” project.

## DATA AVAILABILITY STATEMENT

The data that support the findings of this study are openly available in Human Connectome Project database at https://db.humanconnectome.org/.

The code used for the analysis is available on the GitHub page of the Imaging Language Group: https://github.com/AaltoImagingLanguage/Karhula2026.

## Footnotes

1 Note that aside the model description, which follows nomenclature used in Haakana et al. (2024), N refers to number of subjects, again following typical nomenclature in literature.

2 Hyperparameters *a_σ_* and *b_σ_* are used to define the prior for the noise parameters (*σ* ^−2^) (Gillberg et al., 2016).

