## Supplementary Material for "Machine learning on magnetoencephalography data yields generalizable low-dimensional neural fingerprints that distinguish individuals across task conditions"

#### TABLES

**Table 1.** Mean lnBRRR accuracies (SEM) and corrected p-values for models trained with N=20-44

| Metric | N | Training set | Test set | p (corrected) |
|---|---|---|---|---|
| PLV | 20 | 0.99 (0.013) | 0.68 (0.002) | 0.002 |
| | 25 | 0.97 (0.030) | 0.68 (0.004) | 0.002 |
| | 30 | 0.94 (0.025) | 0.75 (0.020) | 0.002 |
| | 35 | 0.95 (0.021) | 0.73 (0.003) | 0.002 |
| | 44 | 0.94 (0.017) | 0.74 (0.003) | 0.002 |
| PSD | 20 | 0.89 (0.087) | 0.76 (0.042) | 0.002 |
| | 25 | 0.87 (0.030) | 0.78 (0.054) | 0.002 |
| | 30 | 0.83 (0.025) | 0.78 (0.054) | 0.002 |
| | 35 | 0.80 (0.029) | 0.78 (0.020) | 0.002 |
| | 44 | 0.81 (0.065) | 0.76 (0.043) | 0.002 |

† Training set random chance = ∼0.05-0.02 (N=20-44)

† Test set random chance = ∼0.02

**Table 2.** Mean accuracies (SEM) on the training set with full data and latent spaces. Full data identification was performed with Pearson's correlation (Correlation) and cosine distance (Cos dist.) as the distance function. In the PCA, LDA, and lnBRRR latent spaces, the identification was performed with cosine distance as the distance function.

| | Full data | | Latent space | | |
|---|---|---|---|---|---|
| Metric | Correlation | Cos dist. | PCA | LDA | LnBRRR |
| PSD | 0.93 (0.007) | 0.87 (0.010) | 0.79 (0.012) | **0.90 (0.009)** | 0.81 (0.012) |
| PLV ($\delta$) | 0.95 (0.009) | 0.95 (0.007) | 0.70 (0.020) | 0.02 (0.004) | **0.73 (0.013)** |
| PLV ($\theta$) | 0.98 (0.007) | 0.97 (0.005) | **0.86 (0.021)** | 0.08 (0.005) | **0.86 (0.011)** |
| PLV ($\alpha$) | 0.98 (0.003) | 0.97 (0.006) | 0.86 (0.019) | 0.17 (0.020) | **0.87 (0.012)** |
| PLV (low $\beta$) | 0.98 (0.002) | 0.98 (0.004) | **0.92 (0.008)** | 0.22 (0.019) | 0.91 (0.012) |
| PLV (high $\beta$) | 0.99 (0.004) | 0.99 (0.004) | **0.94 (0.007)** | 0.29 (0.013) | **0.94 (0.005)** |
| PLV ($\gamma$) | 1.0 (0) | 1.0 (0) | **0.94 (0.013)** | 0.33 (0.027) | **0.94 (0.012)** |
| iPLV ($\delta$) | 0.14 (0.009) | 0.10 (0.008) | 0.04 (0.009) | 0.0 (0) | **0.05 (0.009)** |
| iPLV ($\theta$) | 0.17 (0.006) | 0.19 (0.011) | **0.11 (0.011)** | 0.00 (0.004) | 0.10 (0.012) |
| iPLV ($\alpha$) | 0.69 (0.013) | 0.50 (0.007) | **0.45 (0.015)** | 0.07 (0.007) | 0.39 (0.013) |
| iPLV (low $\beta$) | 0.18 (0.013) | 0.16 (0.006) | **0.14 (0.008)** | 0.01 (0.004) | 0.10 (0.007) |
| iPLV (high $\beta$) | 0.24 (0.015) | 0.24 (0.013) | 0.15 (0.012) | 0.0 (0) | **0.16 (0.013)** |
| iPLV ($\gamma$) | 0.13 (0.005) | 0.13 (0.008) | 0.06 (0.011) | 0.0 (0) | **0.07 (0.009)** |
| PLM ($\delta$) | 0.05 (0.006) | 0.04 (0.004) | **0.02 (0.005)** | **0.02 (0.004)** | 0.01 (0.004) |
| PLM ($\theta$) | 0.71 (0.011) | 0.47 (0.016) | **0.31 (0.013)** | 0.01 (0.005) | 0.25 (0.023) |
| PLM ($\alpha$) | 0.92 (0.005) | 0.69 (0.007) | **0.65 (0.016)** | 0.26 (0.018) | 0.48 (0.030) |
| PLM (low $\beta$) | 0.96 (0.005) | 0.85 (0.015) | **0.78 (0.019)** | 0.10 (0.011) | 0.65 (0.020) |
| PLM (high $\beta$) | 0.99 (0.004) | 0.96 (0.004) | **0.79 (0.011)** | 0.08 (0.013) | **0.79 (0.013)** |
| PLM ($\gamma$) | 0.99 (0.002) | 0.98 (0.004) | **0.91 (0.011)** | 0.10 (0.011) | **0.91 (0.011)** |
| AEC ($\delta$) | 0.86 (0.004) | 0.79 (0.014) | 0.38 (0.014) | 0.01 (0.004) | **0.43 (0.013)** |
| AEC ($\theta$) | 0.94 (0.007) | 0.79 (0.009) | **0.60 (0.017)** | 0.02 (0.004) | 0.53 (0.021) |
| AEC ($\alpha$) | 0.90 (0.010) | 0.76 (0.008) | **0.58 (0.005)** | 0.14 (0.010) | 0.51 (0.018) |
| AEC (low $\beta$) | 0.98 (0.002) | 0.78 (0.010) | **0.68 (0.012)** | 0.15 (0.015) | 0.55 (0.015) |
| AEC (high $\beta$) | 0.98 (0.006) | 0.89 (0.010) | **0.75 (0.015)** | 0.15 (0.012) | 0.64 (0.011) |
| AEC ($\gamma$) | 0.98 (0.004) | 0.98 (0.004) | 0.85 (0.018) | 0.11 (0.012) | **0.86 (0.010)** |

$^{\dagger}$ Training set N=54-55, random chance = $\sim$0.02

**Table 3.** Mean accuracies (SEM) for PLV data with LDA decision function and cosine distance in LDA latent space

| Metric | LDA | LDA (cos dist.) |
|---|---|---|
| PLV ($\delta$) | 0.83 (0.007) | 0.02 (0.004) |
| PLV ($\theta$) | 0.91 (0.012) | 0.08 (0.005) |
| PLV ($\alpha$) | 0.97 (0.011) | 0.17 (0.020) |
| PLV (low $\beta$) | 0.97 (0.005) | 0.22 (0.019) |
| PLV (high $\beta$) | 0.97 (0.007) | 0.29 (0.013) |
| PLV ($\gamma$) | 0.99 (0.005) | 0.33 (0.027) |

**Table 4.** Mean lnBRRR accuracies (SEM) for task data

| Task | Metric | Training set | Test set |
|---|---|---|---|
| WM | PSD | 0.93 (0.015) | 0.93 (0.017) |
| | PLV ($\delta$) | 0.55 (0.021) | 0.43 (0.089) |
| | PLV ($\theta$) | 0.78 (0.011) | 0.55 (0.072) |
| | PLV ($\alpha$) | 0.86 (0.019) | 0.77 (0.028) |
| | PLV (low $\beta$) | 0.95 (0.005) | 0.76 (0.030) |
| | PLV (high $\beta$) | 0.94 (0.011) | 0.92 (0.031) |
| | PLV ($\gamma$) | 0.96 (0.004) | 0.91 (0.028) |
| Story | PSD | 0.85 (0.019) | 0.86 (0.048) |
| | PLV ($\delta$) | 0.84 (0.027) | 0.77 (0.062) |
| | PLV ($\theta$) | 0.90 (0.015) | 0.80 (0.065) |
| | PLV ($\alpha$) | 0.94 (0.006) | 0.87 (0.024) |
| | PLV (low $\beta$) | 0.89 (0.009) | 0.79 (0.025) |
| | PLV (high $\beta$) | 0.92 (0.014) | 0.84 (0.042) |
| | PLV ($\gamma$) | 0.92 (0.011) | 0.81 (0.025) |
| Math | PSD | 0.75 (0.019) | 0.77 (0.041) |
| | PLV ($\delta$) | 0.76 (0.014) | 0.67 (0.089) |
| | PLV ($\theta$) | 0.80 (0.019) | 0.69 (0.076) |
| | PLV ($\alpha$) | 0.85 (0.018) | 0.81 (0.084) |
| | PLV (low $\beta$) | 0.85 (0.013) | 0.72 (0.046) |
| | PLV (high $\beta$) | 0.83 (0.020) | 0.75 (0.050) |
| | PLV ($\gamma$) | 0.85 (0.017) | 0.68 (0.084) |

† Training set N=54-55, random chance = ∼0.02

† Test set N=13-14, random chance = ∼0.07

**Table 5.** Mean lnBRRR accuracies (SEM) for identification with resting-state data and models from each condition

| | | Training set | Test set |
|---|---|---|---|
| Model | Metric | lnBRRR (SEM) | lnBRRR (SEM) |
| RS | Power spectra | 0.81 (0.012) | 0.83 (0.035) |
| | PLV ($\delta$) | 0.73 (0.013) | 0.62 (0.050) |
| | PLV ($\theta$) | 0.86 (0.011) | 0.72 (0.026) |
| | PLV ($\alpha$) | 0.87 (0.012) | 0.81 (0.029) |
| | PLV (low $\beta$) | 0.91 (0.012) | 0.87 (0.047) |
| | PLV (high $\beta$) | 0.94 (0.005) | 0.83 (0.017) |
| | PLV ($\gamma$) | 0.94 (0.012) | 0.85 (0.027) |
| WM | Power spectra | 0.79 (0.020) | 0.81 (0.041) |
| | PLV ($\delta$) | 0.46 (0.019) | 0.56 (0.054) |
| | PLV ($\theta$) | 0.67 (0.013) | 0.66 (0.034) |
| | PLV ($\alpha$) | 0.73 (0.014) | 0.73 (0.045) |
| | PLV (low $\beta$) | 0.88 (0.006) | 0.78 (0.090) |
| | PLV (high $\beta$) | 0.86 (0.010) | 0.87 (0.053) |
| | PLV ($\gamma$) | 0.89 (0.015) | 0.85 (0.064) |
| Story | Power spectra | 0.78 (0.019) | 0.81 (0.027) |
| | PLV ($\delta$) | 0.44 (0.018) | 0.57 (0.069) |
| | PLV ($\theta$) | 0.65 (0.019) | 0.62 (0.021) |
| | PLV ($\alpha$) | 0.75 (0.014) | 0.80 (0.041) |
| | PLV (low $\beta$) | 0.86 (0.024) | 0.87 (0.038) |
| | PLV (high $\beta$) | 0.83 (0.010) | 0.81 (0.035) |
| | PLV ($\gamma$) | 0.85 (0.010) | 0.83 (0.037) |
| Math | Power spectra | 0.72 (0.014) | 0.73 (0.028) |
| | PLV ($\delta$) | 0.48 (0.012) | 0.54 (0.074) |
| | PLV ($\theta$) | 0.69 (0.013) | 0.61 (0.030) |
| | PLV ($\alpha$) | 0.74 (0.017) | 0.80 (0.038) |
| | PLV (low $\beta$) | 0.86 (0.024) | 0.84 (0.047) |
| | PLV (high $\beta$) | 0.81 (0.025) | 0.87 (0.038) |
| | PLV ($\gamma$) | 0.85 (0.009) | 0.83 (0.034) |

### FIGURES

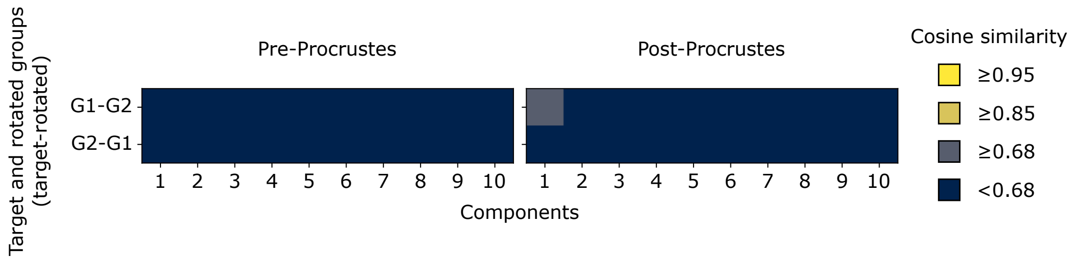


**Figure 1.** Resting-state gamma-band PLV lnBRRR $\Gamma$ similarities between training samples. Cosine similarities between orthonormalized $\Gamma$ matrix rows before and after orthogonal Procrustes analysis were used to assess whether resting-state lnBRRR solutions derived from two equal-sized training sets are similar. X-axis indicates the $\Gamma$ rows (Components) and y-axis indicates the target and rotated groups (G1=group 1, G2=group 2). Cosine similarity was divided into four classes: practically equal ($\geq 0.95$), similar ($\geq 0.85$), poorly similar ($\geq 0.68$), and no similarity ($< 0.68$).

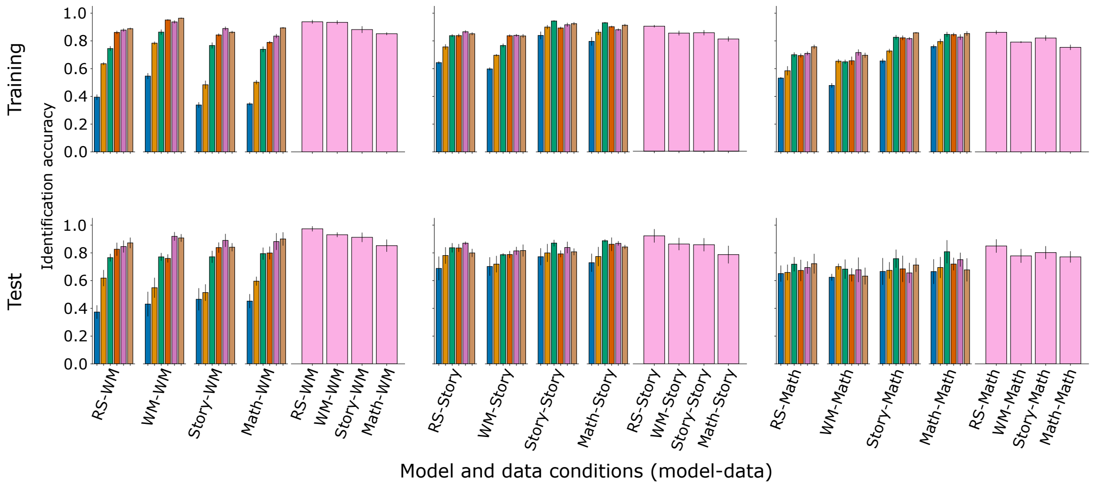


**Figure 2.** Cross-condition lnBRRR accuracies with task condition data. PLV and power spectra data, and lnBRRR models trained on data from each condition were used to assess whether lnBRRR models trained on task data can identify individuals based on their resting-state data. Mean identification accuracies over the CV folds and SEM are shown for the training (upper row) and test (lower row) sets. The data condition is shown above the plots and the task condition of the model used for identification is shown on the x-axis. The original model (e.g. WM model for WM data) results are shown for comparison.

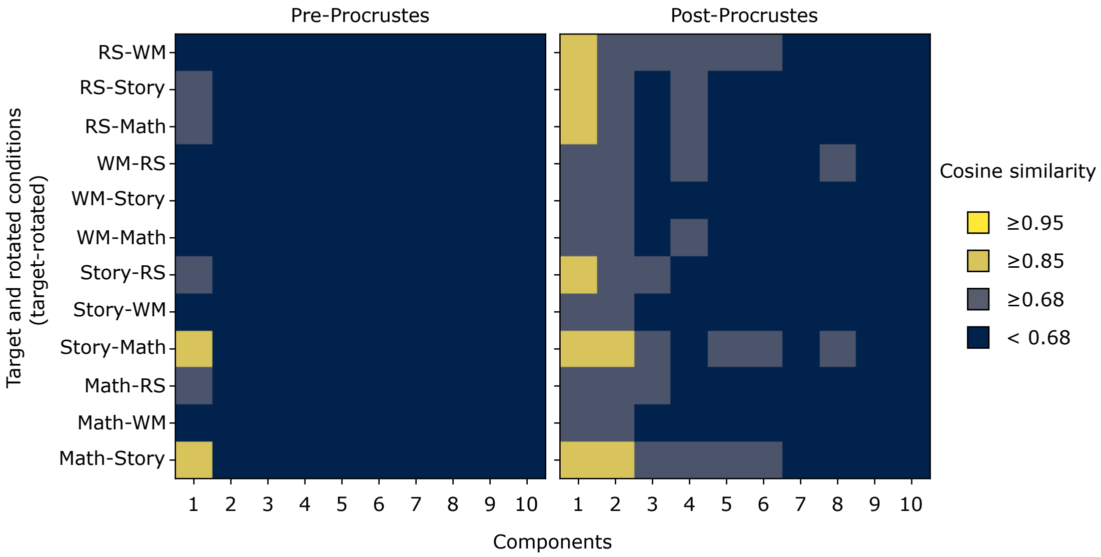


**Figure 3.** Between-task $\Gamma$ similarities for gamma-band PLV lnBRRR models. The similarity between lnBRRR solutions from different conditions was measured by cosine similarity between orthonormalized $\Gamma$ matrix rows. Similarity was measured before and after rotational alignment by Orthogonal Procrustes analysis. X-axis indicates the $\Gamma$ rows (Components) and y-axis indicates the condition pair examined. Cosine similarity was divided into four classes: practically equal ($\geq 0.95$), similar ($\geq 0.85$), poorly similar ($\geq 0.68$), and no similarity ($< 0.68$).

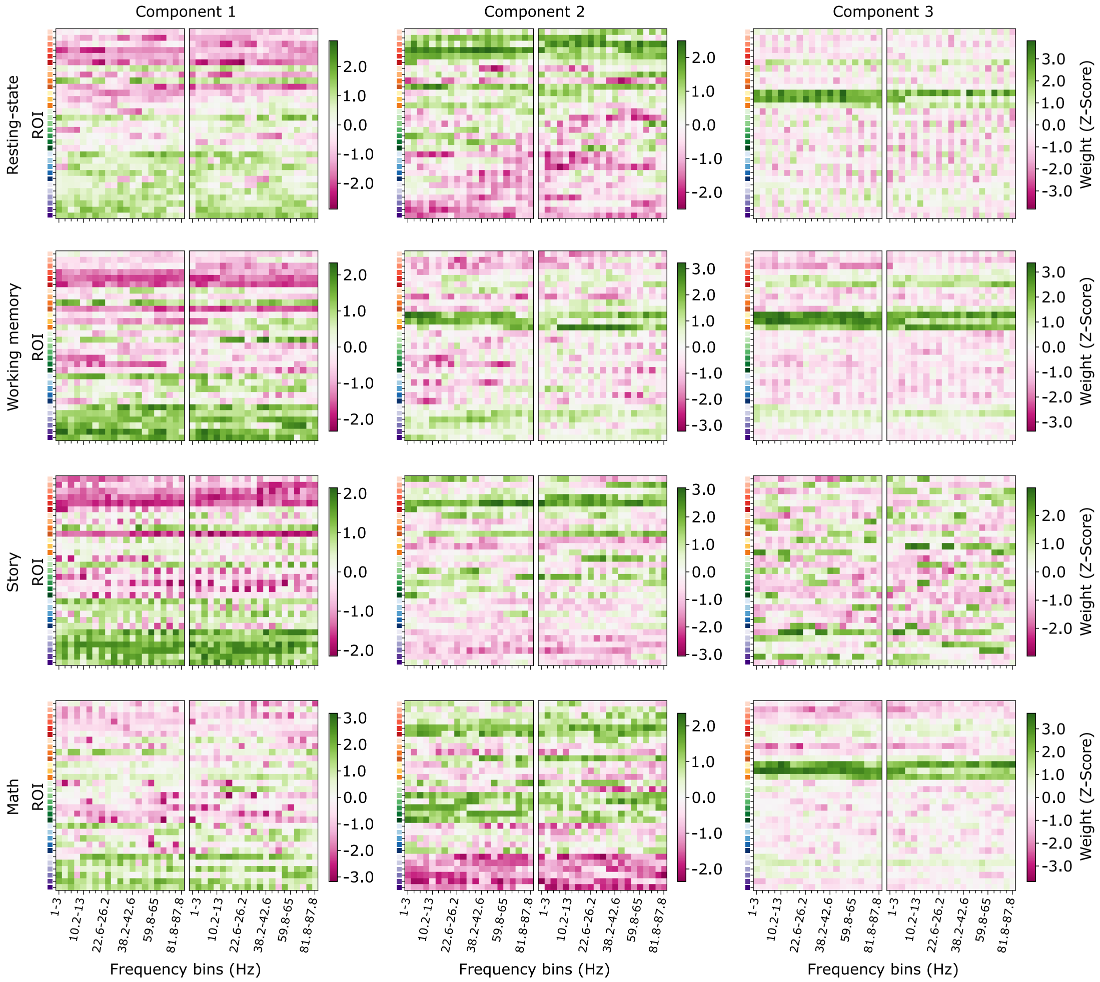


**Figure 4.** All power spectral lnBRRR weights. LnBRRR was trained on resting-state, working-memory, story, and math task power spectra from the same 68 participants. All the weights for the first three unrotated components are shown for each model. Note that the colorbars have different scales between components and conditions.
